# Local fluidization of an active cytoplasmic gel partitions large cells

**DOI:** 10.64898/2026.08.11.744254

**Authors:** Luolan Bai, Christine M. Field, Ai Kiyomitsu, Yunyi Shen, Natalia D. Orlovsky, Tomomi Kiyomitsu, Timothy J. Mitchison

**Affiliations:** Department of Systems Biology, Harvard Medical School, Boston, MA, USA; Department of Molecular and Cellular Biology, Harvard University, Cambridge, MA, USA; Okinawa Institute of Science and Technology Graduate University, 1919-1 Tancha, Onna-son, Kunigami-gun, Okinawa, 904-0495, Japan; Department of Electrical Engineering and Computer Science, Massachusetts Institute of Technology, Cambridge, MA, USA

## Abstract

Early animal embryos undergo rapid cleavages that partition cytoplasmic volumes orders of magnitude larger than those of somatic cells^1^. Each division must reposition nuclei and centrosomes and distribute organelles within minutes, over distances up to hundreds of micrometers^2^. Cleavage furrows are positioned by microtubule asters^3,4^, but the mechanical mechanism for long-range transport of cytoplasmic components before cytokinesis was unknown. Here, we show that cytoplasm behaves as a locally switchable active material. Fluidization at the midplane allows bulk actomyosin to convert a local mechanical asymmetry into directed global flows of all components as a composite material. Using an actin-intact cycling *Xenopus* egg extract together with *Xenopus* and medaka embryos, we find that F-actin mechanically couples microtubule asters, organelles, nuclei and centrosomes into a gel-like composite that propagates forces over hundreds of micrometers. After mitosis, Aurora B kinase patterns a locally fluidized midplane, from which myosin-II contractility drives coherent cytoplasmic flows. A fluid dynamics model accounts for the observed flow geometry and rates. Our results reveal how local control of the material state of cytoplasm converts mitotic symmetry breaking into long-range intracellular transport and identify bulk actomyosin as the active stress generator that partitions embryonic cytoplasm as a composite gel.

## Main

Achieving spatial organization across length scales is a fundamental challenge in biology. This challenge is especially acute in early animal embryos, which cleave rapidly without growth and must partition cytoplasmic volumes 10^3^-to 10^5^-fold larger than those of somatic cells^1^. In large embryos, nuclei, centrosomes, organelles and cytoskeletal networks are redistributed over hundreds of micrometers within minutes. For example, in the *Xenopus* zygote, centrosomes and chromatin move more than 100 µm away from the midplane between anaphase and cytokinesis within ∼10 minutes^2^. How intracellular contents are transported over such distances remains poorly understood.

Mitotic mechanics is usually understood in terms of molecular machines acting on discrete cellular structures. After each mitosis, spindle poles nucleate microtubule asters that position nuclei and specify the cleavage plane^5–7^. Organelle partitioning is similarly often framed through cargo–motor interactions along cytoskeletal tracks^8^. These local mechanisms are powerful in small cells, where spindle size, microtubule length and cell radius are comparable. In large embryonic cells, however, cytoplasmic organization must occur over length scales far exceeding individual microtubules and the mitotic spindle^3^. This raises the question of whether local microtubule- and motor-based mechanisms are sufficient to move bulk cytoplasm coherently over hundreds of micrometers.

An alternative view is that cytoplasm itself can act as a mechanically integrated material. Cytosol, organelles, cytoskeletal polymers and motors together form a composite viscoelastic medium whose mechanical properties depend on composition, length scale and timescale^9–11^. At scales larger than the cytoskeletal mesh size, filament networks with associated motors can behave as active materials that transmit stress and generate flows^12–17^. The large size of early embryos also decreases the surface-to-volume ratio, likely increasing the relative mechanical contribution of bulk cytoplasm compared with the cortex. Although cytoplasmic material properties have been measured in many systems^18–20^, whether cells actively pattern these properties to drive large-scale intracellular transport during division remains unclear.

Here we show that bulk cytoplasm behaves as a locally switchable active material for post-mitotic organization. Using an actin-intact cycling *Xenopus* egg extract together with biochemical and direct physical perturbations, embryo imaging and fluid dynamics modeling, we find that F-actin mechanically couples microtubule asters, organelles, nuclei and centrosomes into a gel-like composite that stabilizes cytoplasmic organization and transmits force over hundreds of micrometers. After mitosis, Aurora B kinase patterns a locally fluidized midplane, and myosin-II contractility drives coherent flows that move cytoplasmic components away from the midplane. A minimal model shows that contractile stress acting in a cytoplasm with local mechanical asymmetry is sufficient to drive cytoplasmic partitioning.

### Cortex-free system for investigating mechanics of early embryo partitioning

Studying intracellular mechanics in dividing embryos is complicated by concurrent mechanical inputs from the cell surface and bulk cytoplasm. To investigate cytoplasmic mechanics independent of the cortex, we established an egg extract system that faithfully reconstitutes the physical organization of embryonic cytoplasm. Standard cycling *Xenopus laevis* egg extracts have revealed how microtubule asters self-organize the geometry of cytoplasmic compartments^21,22^, but they face two limitations for studying post-mitotic cytoplasmic mechanics: (1) F-actin is chemically disrupted during preparation, removing the mechanical contribution of at least one major cytoskeletal network^21–23^; (2) mitotic partitioning of cytoplasmic contents in these extracts is often unstable, lacking the accurate and persistent nuclear and centrosomal separation observed in embryos^22,24^. We therefore established an actin-intact cycling egg extract that faithfully reconstituted cytoplasmic mechanics, supported repeated cell-free division of cytoplasmic compartments, enabled high-resolution long-term live imaging with limited global contraction^25^, and allowed direct mechanical perturbation (Fig. 1a–d, Supplementary Videos 1 and 2, and Methods). Sperm nuclei with attached centrosomes added to the extract duplicated and segregated once per cycle (Fig. 1b), with efficient chromatin segregation (Extended Data Fig. 1a). After each mitosis the cell-like compartments partitioned in two as daughter centrosomes and nuclei moved hundreds of µm apart and centered (Fig. 1b), confirming that long-range positioning does not require interaction between microtubules and the cell cortex, consistent with our previous proposal^2,26^. Robust post-mitotic separation between centrosomes and asters progressed for hours over repeated cycles (Fig. 1c). F-actin and keratin both organized into fibrous cytoplasmic meshworks (Fig. 1d), confirming that all three cytoskeletal systems were reconstituted. To our knowledge, this is the most physiological cell-free reconstitution of embryonic cytoplasm reported to date.

**Fig. 1:**
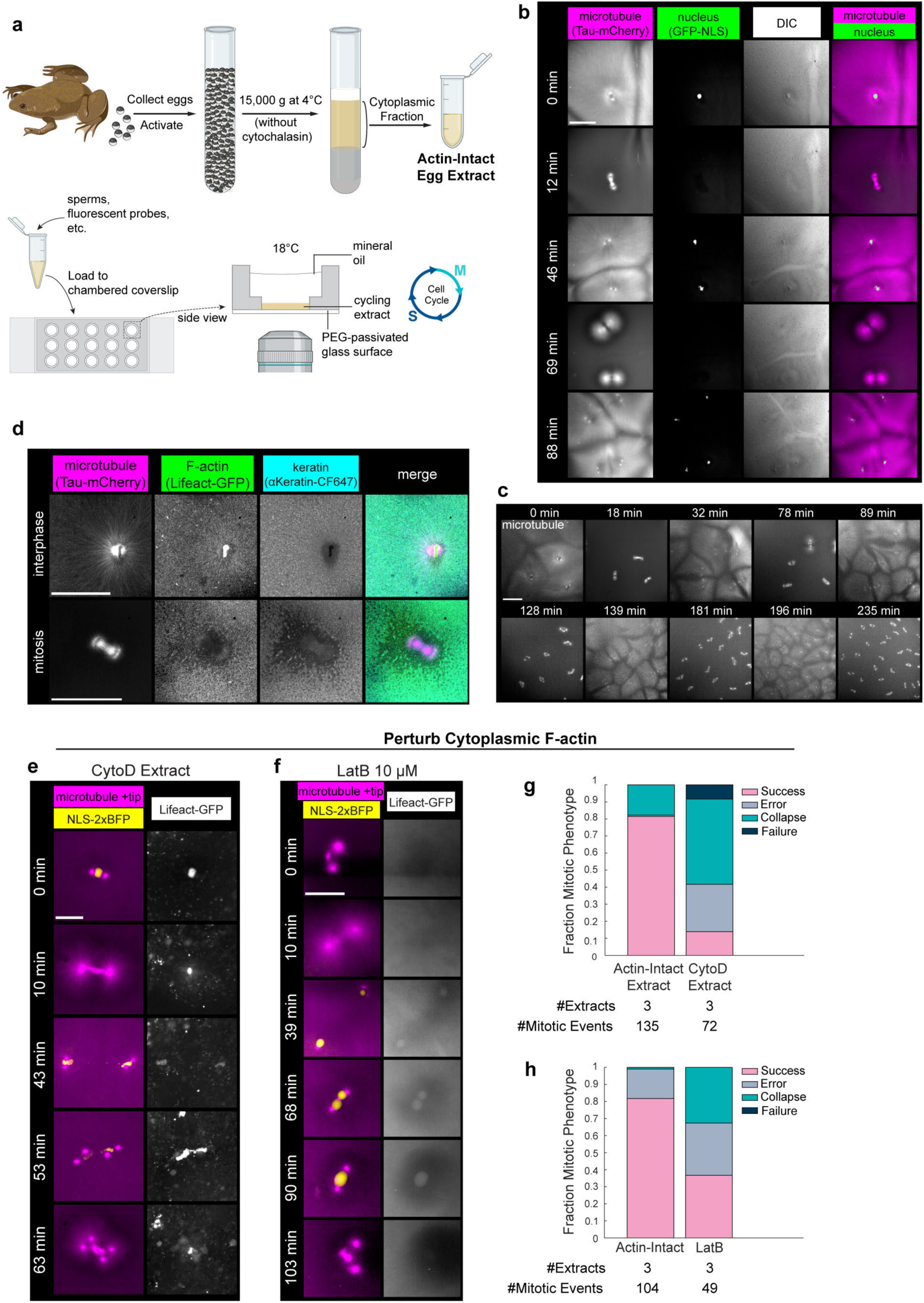
Cytoplasmic F-actin enables robust post-mitotic separation in cell-free extract. **a**, Schematic overview of the procedure to prepare and image actin-intact cycling egg extract. **b**, Microtubule and nuclear dynamics during cell cycle progression of the egg extract alternating between interphase and mitosis, visualized with Tau-mCherry and GFP-NLS. Images are maximum intensity projection (MaxIP) from 2 Z-stacks 10 µm apart. **c**, Time lapse of continuous cell cycle progression over 4 hours. Microtubules visualized with Tau-mCherry. **d**, Confocal images of microtubules (Tau-mCherry), F-actin (Lifeact-GFP) and keratin filaments (anti-keratin labelled with CF647 dye) during interphase and mitosis in cycling extract. **e**, Daughter centrosomes and nuclei collapsing back (referred to as “Collapse” in **g**) after initial separation, in cycling egg extract prepared with cytochalasin D (CytoD extract). Microtubules and centrosomes visualized with EB1-mApple, which binds to microtubule plus ends. Nuclei visualized with NLS-2xBFP. Actin fragments and aggregates from CytoD treatment visualized with Lifeact-GFP. **f**, Daughter centrosomes and nuclei collapsing back (“Collapse” in **h**) after initial separation, in cycling egg extract treated with 10 µM latrunculin B (LatB). **g**, Mitotic outcomes among 3 pairs of actin-intact extract and CytoD extract. **h**, Mitotic outcomes among 3 pairs of actin-intact extracts, untreated versus treated with 50 µM LatB. Scale bars, 200 µm for **b**, **c**, **d**; 100 µm for **e**, **f**.

### F-actin facilitates robust post-mitotic separation

Actin-perturbing drugs were used to test if F-actin facilitates robust segregation of nuclei and partitioning of cytoplasm (Fig. 1e–h). Actin-intact extract and a cytochalasin D–treated counterpart (“CytoD extract”) were prepared in parallel from the same batches of eggs, and their mitotic outcomes were scored (Fig. 1e, g, Extended Data Fig. 1b, and Methods). CytoD extracts exhibited multifaceted defects: increased errors in chromosome and centrosome segregation (“Error”), and even when segregation initially succeeded, unstable post-mitotic separation in which daughter nuclei and centrosomes collapsed back together and formed multipolar spindles in the next cycle (“Collapse”, Fig. 1e). Across three paired preparations, actin-intact extracts succeeded in 81% of divisions versus 14% for CytoD extracts, with Collapse being the most frequent outcome (50%) (Fig. 1g). Whereas CytoD, a barbed-end capping drug, replaces the meshwork with fragments and aggregates, latrunculin B (LatB) sequesters actin monomers and abolishes filament assembly^27^. LatB likewise increased both Collapse and Error (Fig. 1f, h), confirming that the defects reflect loss of the F-actin network rather than being cytochalasin-specific. We concluded that an intact cytoplasmic F-actin network is required for robust segregation and stable post-mitotic partitioning of nuclei and centrosomes. Our observation on early segregation defects is consistent with a role of nuclear and spindle actin in meiotic and mitotic fidelity^28–31^, so we instead focused on how cytoplasmic F-actin contributes to long-distance transport and partitioning after mitosis.

### Coordinated partitioning of cytoplasm after mitosis

We next investigated how other components of cytoplasm are partitioned after mitosis. After anaphase, the sister asters grow until their microtubules meet at the midplane (Fig. 2a), where antiparallel bundles recruit the chromosomal passenger complex (CPC), which blocks microtubule interpenetration and specifies the cleavage plane^3,4^. F-actin and keratin networks were present throughout the cytoplasm in interphase and then reorganized at the midplane after anaphase (Fig. 2a). Individual filaments were not resolved at the low-magnification imaging required to measure global behaviors. After mitotic onset, F-actin persisted only at low density in spindles (Fig. 1d, bottom row and Fig. 2a, 31 min), while the keratin meshwork disassembled throughout the cytoplasm (Fig. 2a, 23–31 min). From late anaphase into the following interphase, a gap in F-actin density developed at the midplane and widened as the asters grew, exceeding 100 µm by interphase; as keratin reassembled, it too re-organized around the midplane, leaving a thin layer of keratin aggregates at its center.

**Fig. 2:**
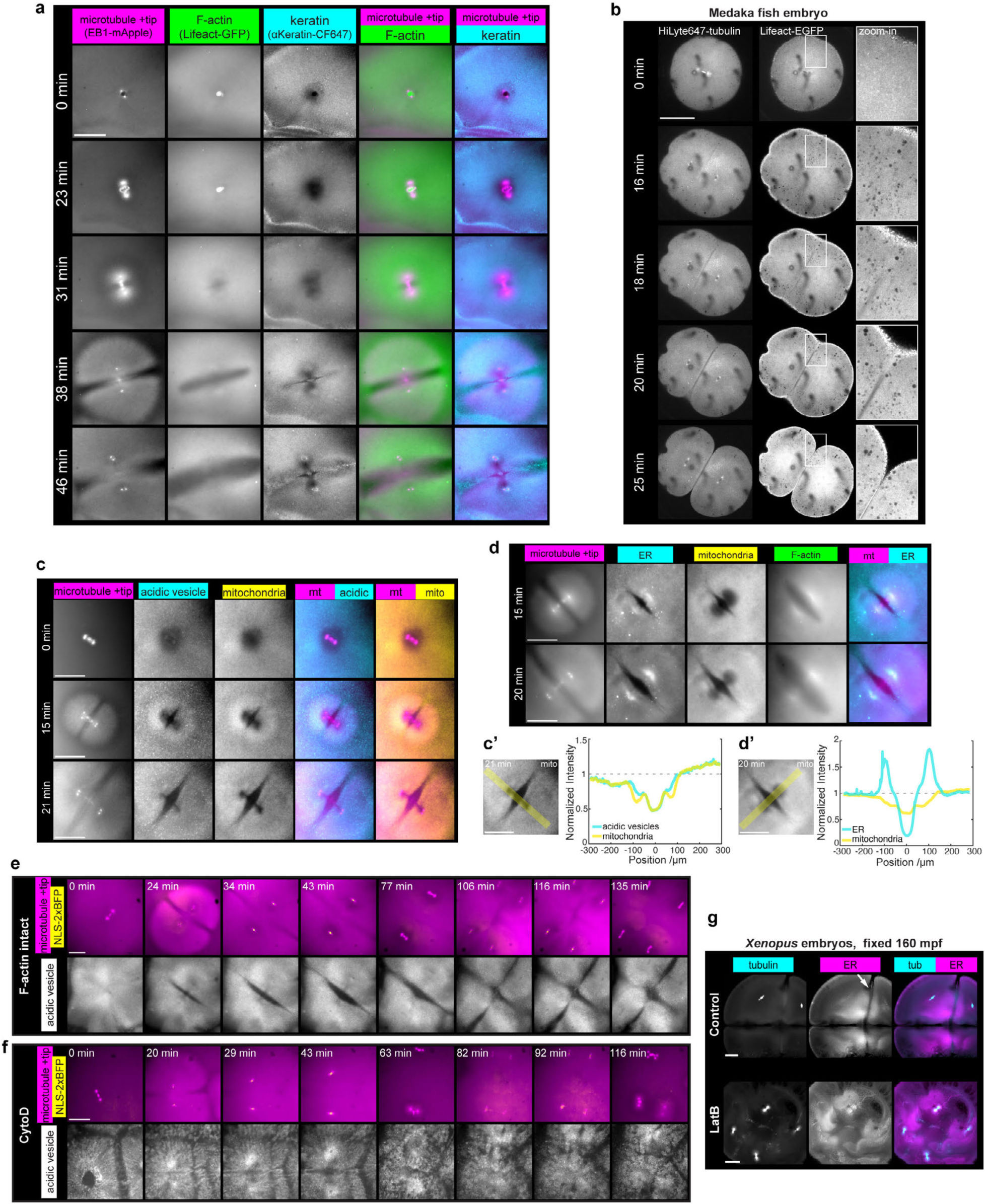
Coordinated partitioning of cytoskeletal and organelle networks after mitosis is supported by F-actin. **a**, Time lapse of microtubules, F-actin and keratin filaments across mitotic progression in actin-intact cycling extract. **b**, Live images showing microtubules (HiLyte647-tubulin, injected) and F-actin (Lifeact-EGFP) in a cleaving medaka zygote. Zoom-in: F-actin depletion at the midplane. **c** and **d**, Live images of acidic vesicles (LysoTracker Deep Red), ER (Vybrant DiD) and mitochondria (MitoView 405) in actin-intact cycling egg extract. **c’** and **d’**, Organelle density quantified for **c** and **d** along the axis of separation, averaged over a 50-µm-wide band spanning centrosome pairs (yellow shaded in the mitochondria image). **e** and **f**, Time lapse over 2 hours of microtubules (EB1-mApple), nuclei (NLS-2xBFP) and acidic vesicles (Lysotracker Deep Red) in cycling egg extracts with intact F-actin (**e**) or prepared with CytoD (**f**). Note the variable distribution of acidic vesicles in CytoD extract both spatially and temporally, including poor or unstable partitioning at the midplane. **g**, Immunofluorescence images of *Xenopus laevis* embryos fixed at 160-minutes post-fertilization (mpf) and viewed from the animal pole, visualizing microtubules (anti-tubulin-AF647) and ER (anti-cytLnp-AF568). Top: untreated, showing maximum intensity projection (MaxIP) from 3 Z-stacks. White arrow: ER depletion beneath an upcoming furrow. Bottom: treated with 10 µg/ml (25 µM) LatB from 20 mpf, showing MaxIP from 4 Z-stacks. All scale bars, 200 µm. For **c**, **d**, **e** and **f**, time 0 indicates the first frame with assembled spindles (and fully disassembled nuclei).

In live medaka embryos post-fertilization cytoplasm occupies a ∼500-µm-wide, ∼100-µm-thick blastodisc^6,32^. In this system, a sharp zone of F-actin depletion formed between asters at the midplane, deep beneath the actin-rich cortex (Fig. 2b, 18-20 min). The furrow subsequently ingressed precisely where F-actin had cleared (Fig. 2b, 25 min); similar clearing was recently reported in zebrafish^33^. These observations show that our extract system faithfully recapitulates live embryo dynamics.

We next imaged the three major organelle populations of egg extract, including acidic vesicles (lysosomes and endosomes), mitochondria, and endoplasmic reticulum (ER) (Fig. 2c–d). As sister centrosomes separated after anaphase, all three became progressively depleted from the midplane, generating organelle-free zones between the asters (Fig. 2c–d and Supplementary Video 3). Although each organelle displayed a distinct distribution near the centrosomes, all showed strong and consistent depletion at the midplane (Fig. 2c′, 2d′). The clearing of biochemically and functionally unrelated organelles from the same zone argues that partitioning is driven by a global reorganization of bulk cytoplasm rather than by organelle-specific motor transport, paralleling the local clearing of cytoskeleton. This depletion was often stable beyond the following mitosis, which further sub-partitioned the cytoplasm (Fig. 2e and Supplementary Video 3). Organelles also partitioned along the midplane between mitosis and cytokinesis in live medaka and fixed *Xenopus* embryos as illustrated by ER distribution (Extended Data Fig. 2a, a’, b, c and Fig. 2g top row). We conclude that coordinated global partitioning of cytoplasmic components occurs independent of, and prior to, cytokinesis in cell-free extract and large embryos. That all cytoskeletal and organelle networks were remodeled to lower density along the midplane suggests a local switch-like transition of the cytoplasm as a material.

### Coordinated partitioning requires cytoplasmic F-actin

Disrupting F-actin with CytoD or LatB led to uneven and chaotic distribution of acidic vesicles with aggregation around centrosomes and frequent large-scale flows, in contrast to their smooth, stable and largely uniform distribution in actin-intact extract apart from sharp midplane depletion (Fig. 2e–f, Extended Data Fig. 2d-f and Supplementary Video 4). Vesicle partitioning was less efficient and more variable, showing weak or late clearance and re-fusion after initial separation, and these defects often coincided with the collapse of centrosomes and nuclei (Fig. 2f, Extended Data Fig. 2d, e, e’, f, f’ and Supplementary Video 4). The shift of acidic vesicle density correlated with centrosome movement (Extended Data Fig. 2e’, f’), consistent with the proposed co-movement between organelles and asters^26^. Similar defects could be noticed in recent studies conducted with standard cytochalasin-treated cycling egg extracts^22^. In fixed *Xenopus* embryos, ER was progressively depleted from the midplane after anaphase and before cytokinesis in untreated embryos, but not in LatB-treated embryos, which failed to divide and showed disorganized ER from the first anaphase onward (Fig. 2g and Extended Data 2b, c, c’). Bulk F-actin is therefore required for the coordinated partitioning of organelles in extract and embryos.

### Aurora B kinase switches cytoplasm to a fluidized state at the midplane

Aurora B kinase as part of the CPC complex is a strong candidate for breaking symmetry at the midplane^4,34^. Across all three systems — cycling extract, *Xenopus* embryos, and medaka embryos — the CPC, marked by INCENP or Aurora B, localized to the midplane between sister asters after anaphase, coincident with the zones from which F-actin and organelles were cleared (Fig. 3a–c, Extended Data Fig. 3a, Supplementary Video 5). We tested the role of Aurora B kinase activity using the small molecule inhibitor barasertib-HQPA. Complete inhibition abolished spindle assembly (Extended Data Fig. 3b); we therefore used partial or timed inhibition, which permitted spindle and aster formation but inhibited subsequent recruitment of the CPC to the midplane. Partial or timed inhibition both prevented the local depletion of microtubules, F-actin and mitochondria at the midplane (Fig. 3c and Extended Data Fig. 3c, d). Aurora B activity is thus required to initiate the midplane clearing of both cytoskeleton and organelles. Initial clearing of both networks likely happens in parallel, as mitochondria were locally depleted around Aurora B in LatB-treated extract (Extended Data Fig. 3e).

**Fig. 3:**
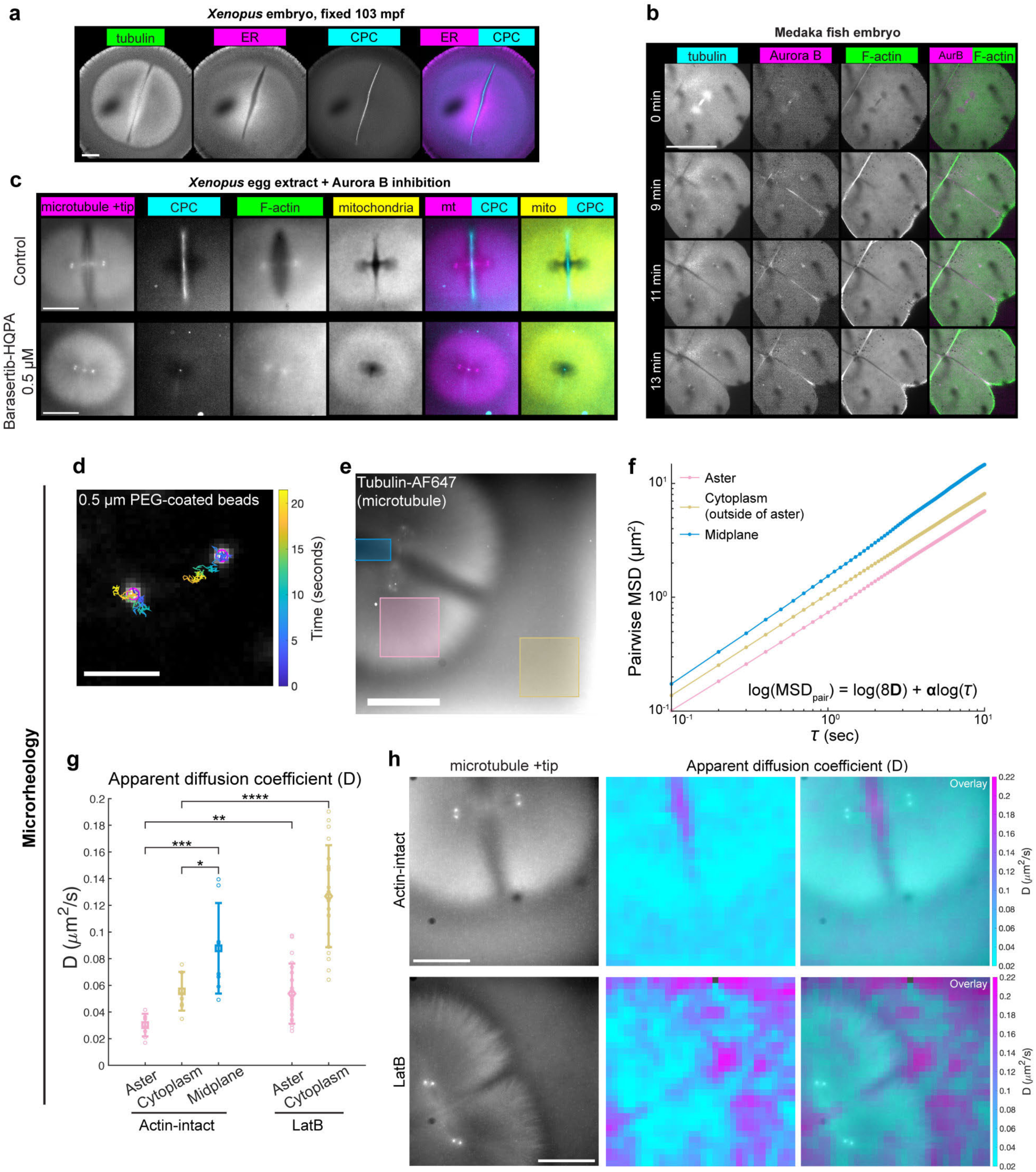
Aurora B kinase patterns a locally fluidized midplane. **a**, Immunofluorescence images of a *Xenopus laevis* zygote fixed after the first mitosis, viewed from the animal pole. CPC (anti-INCENP-AF647) localized to the boundary between sister asters (microtubules imaged with anti-tubulin-AF488) and co-localized with an ER-depleted zone (ER labeled with anti-LNPK-AF568). **b**, Live images of a medaka embryo during 2-cell to 4-cell stage, showing dynamics of microtubules (HiLyte 647-tubulin), CPC (mCherry-Ol-Aurora B) and F-actin (Lifeact-EGFP) from mitosis to cytokinesis. **c**, CPC (anti-INCENP-AF647) localization in *Xenopus* egg extract co-imaged with microtubules (EB1-mApple), F-actin (Lifeact-GFP), and mitochondria (MitoView 405) at late anaphase. Top: untreated. Bottom: treated with 0.5 µM barasertib-HQPA. **d**, Example trajectories of two PEG-coated Fluoresbrite beads of 0.5 µm diameter in particle-tracking microrheology. Scale bar, 5 µm. **e**, Sampled areas in anaphase cycling egg extract with different cytoskeletal composition. **f**, Ensemble-averaged pairwise mean-squared displacement (MSD_pair_) as a function of time lag *τ* for particles within each of the three indicated areas in **e**. **g**, Apparent diffusion coefficient (D) calculated from MSD_pair_ between adjacent beads. For actin-intact controls, 8 areas (65x65 µm^2^) in each category were sampled from independent experiments with 3 different egg extracts. The same extracts were used to prepare latrunculin B-added samples, where 20 areas were sampled in microtubule asters and cytoplasm, respectively. Data are mean ± SD. Also see Extended Data Fig. 4b-e. **h**, Local D calculated within overlapping 42x42 µm^2^ grids, locally averaged, and overlaid onto microtubule images (EB1-mApple). Examples shown are microtubule asters growing against microtubule-free cytoplasm, in actin-intact egg extract (top) or extract treated with 10 µM LatB (bottom). Also see Extended Data Fig. 4f-g. All scale bars except **d**, 200 µm.

We next characterized the physical state change induced by Aurora B kinase activity using particle-tracking microrheology with 0.5 µm PEG-coated beads imaged at 10 Hz. Covalent passivation with PEG prevented protein binding as monitored by prevention of dynein-mediated transport (Extended Data Fig. 4a). We analyzed pairwise mean-squared displacement (MSD_pair_) to correct for bulk drift^35^ (Fig. 3d; Methods). Particle diffusivity correlated with cytoskeletal organization: beads were least diffusive within asters, which contain both microtubules and F-actin (D = 0.0302 ± 0.0085 µm²/s, n = 8), intermediate in microtubule-free bulk cytoplasm which contains F-actin, and most diffusive at the cytoskeleton-and-organelle-depleted midplane (D = 0.0878 ± 0.0340 µm²/s, n = 8) — a roughly three-fold range, with the anomalous exponent approaching free diffusion at the midplane (α = 0.905 ± 0.026, n = 8) (Fig. 3e–h and Extended Data Fig. 4c–g). Depolymerizing F-actin with LatB raised diffusivity throughout the system and reached in cytoplasm an anomalous exponent comparable to the untreated midplane (α = 0.929 ± 0.050, n = 20). The bulk cytoplasm therefore behaves as an F-actin-dependent gel-like composite, within which Aurora B kinase activity carves out a locally fluidized midplane that lacks cytoskeleton and organelles, behaving close to a viscous liquid. By breaking symmetry in this way, Aurora B activity creates a fluidized gap up to ∼50 µm wide within an otherwise gel-like bulk (Fig. 3h, top row). We next asked how this local asymmetry is converted into stable cytoplasmic partitioning over large spatial scales.

### Myosin-II contractility drives aster-spanning flows away from the fluidized midplane

To measure large-scale cytoplasmic motion, we imaged the same PEG-passivated 0.5 µm beads at 10-second intervals (Fig. 4a). Tracking single beads at low time resolution and computing mean velocity over 40 seconds averages out diffusive motion and reports on bulk flow. The beads are approximately the same size as organelles such as mitochondria (Extended Data Fig. 5a), so they report on flow fields that also transport organelles. As the aster pair separated and the fluidized midplane developed, beads throughout each aster moved away from the midplane coherently with one another and with the centrosomes (Fig. 4b, Extended Data Fig. 5b, Supplementary Video 6). Projecting velocities onto the separation axis (*x*) and the midplane (*y*) revealed two velocity plateaus averaging ∼5 µm/min along *x*, each spanning an aster radius, while centrosomes moved at 7.97 ± 2.02 µm/min (n = 19); beads near the midplane flowed inward along *y* (Fig. 4c, d). We hypothesized the inwards flows along *y* serve to balance pressure in an incompressible medium and later confirmed this by modeling (see below). Both the outwards flow along the *x* axis and the inwards flow between asters appeared in live medaka embryos after injecting PEG-coated beads (Extended Data Fig. 5c–h; Supplementary Video 7). Similar flow geometry and rates were reported in a recent study tracking mitochondria in zebrafish embryos, though they were interpreted as evidence of dynein-mediated motility rather than reporting bulk flows^36^. Depolymerizing F-actin destroyed this coherence, removing aster-spanning velocity plateaus and displaying chaotic, variable flows unrelated to aster separation (Extended Data Fig. 5i–m).

**Fig. 4:**
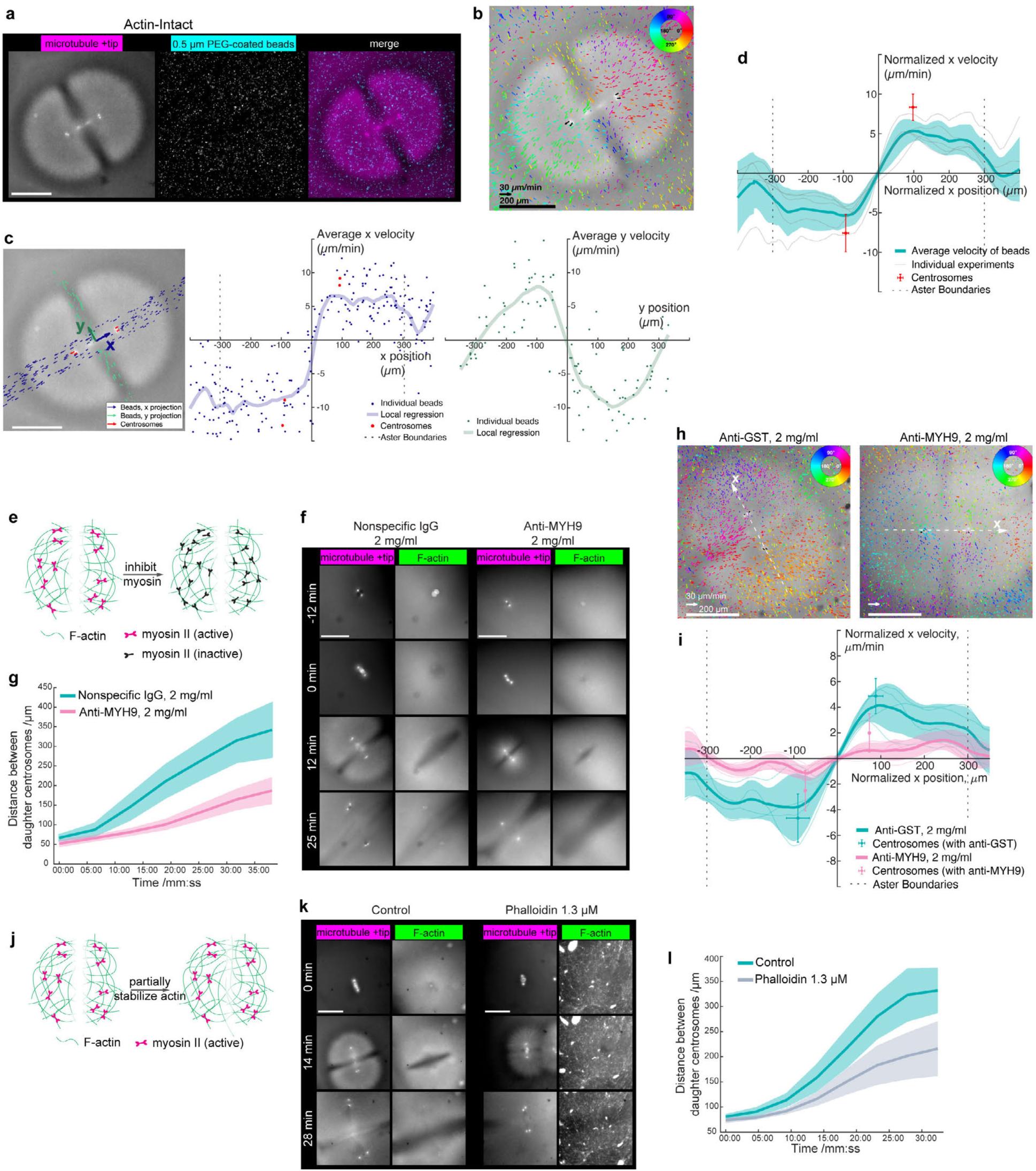
Myosin-II contractility drives aster-spanning cytoplasmic flows away from the fluidized midplane. **a**, Microtubule asters (EB1-mApple) and 0.5 µm PEG-coated Fluoresbrite beads dispersed in cytoplasm, imaged live during late anaphase in actin-intact cycling egg extract. Images are MaxIP from 3 Z-stacks over 10 µm. **b**, Velocity vectors from particle tracking of beads (colored arrows) and centrosomes (black arrows) overlaid on the microtubule image, averaged over 40 seconds. Velocity vectors of beads are color coded by orientation. **c**, Projection of velocity vectors in **b** onto the separation axis (*x* axis) connecting daughter centrosomes, for centrosomes and beads within a 100-µm-wide band along the *x* axis (blue arrows in left panel; middle). Velocity projection of beads along the midplane within a 50-µm-wide band onto the *y* axis (green arrows in left panel; right). *x* and *y* components of particle velocity were plotted against particle position along the *x* and *y* axes, respectively, and local regression with LOWESS was shown. **d**, Ensemble profile of cytoplasmic velocity along the *x* axis averaged from 5 experiments prepared with 5 different actin-intact egg extracts. Asters of similar radius (270 μm – 330 μm) were normalized to 300 μm for aggregated view; bead velocity was normalized so that bulk movement was set to 0. Gray lines show local regression of individual experiments; shaded curve represents mean ± SD of regression curves. **e**, Schematics of myosin inhibition by inhibiting mini-filament formation. **f**, Cytoskeletal dynamics of actin-intact cycling egg extract with 2 mg/ml of nonspecific rabbit IgG and with 2 mg/ml of inhibitory MYH9 antibody. **g**, Quantification of centrosome separation in egg extract with nonspecific rabbit IgG as control and with anti-MYH9. n = 16 aster pairs in control and n = 11 in anti-MYH9-treated extract. See also Extended Data Fig. 6e. **h**, Velocity fields of PEG-passivated 0.5 μm fluorescent beads averaged over 40 seconds, in egg extract with anti-GST as control and with anti-MYH9. Both antibodies were polyclonal and purified from rabbit antiserum. Arrow length shows velocity and color indicates vector orientation. **i**, Ensemble profile of cytoplasmic velocity along the *x* axis in egg extracts with anti-GST and with anti-MYH9, similar as **d**. n = 4 experiments for each group, prepared from 3 different actin-intact egg extracts. Shaded curves represent mean ± SD of regression curves in each group. **j**, Schematics of partially stabilizing F-actin at the midplane. **k**, Cytoskeletal dynamics of untreated and 1.3 µM phalloidin-treated actin-intact cycling egg extract. **l**, Quantification of centrosome separation in control and 1.3 μM phalloidin-treated egg extract. n = 10 aster pairs in control and n = 7 in phalloidin-treated extract. All linear scale bars (except arrows in **b** and **h**), 200 μm. In **d**, **g**, **i** and **l**, data are represented as mean ± SD. For **f**, **g** and **k**, **l**, time 0 = first frame with assembled spindles (and fully disassembled nuclei). Imaging probes include EB1-mApple (microtubule +tip) and Lifeact-GFP (F-actin).

We next asked what powers the flows that transport all components of cytoplasm away from the midplane. Actin depolymerization both fluidizes the cytoplasm and abolishes coherent flows, so actin-perturbing drugs are not a useful test. Indeed, we argue that most studies treating embryos with actin-depolymerizing drugs are hard to interpret because they profoundly change the mechanics of cytoplasm. We instead inhibited myosin-II without perturbing F-actin. Blebbistatin was too insoluble to inhibit myosin-II in our system, so we developed an affinity-purified antibody against the assembly critical domain of non-muscle myosin IIA (MYH9)^37^ based on an earlier study showing antibodies to this region block mini-filament assembly^38^ (Extended Data Fig. 6a–b). Our inhibitory antibody blocked bulk contraction of egg extract (Fig. 4e, Extended Data Fig. 6c–d). Myosin-II inhibition slowed centrosome separation roughly three-fold and slowed the associated bulk flows to a similar extent, without affecting aster growth or midplane actin depletion (Fig. 4f–i, Extended Data Fig. 6e). Contractile stress generated by actomyosin therefore powers the long-range flows that separate asters and carry cytoplasmic contents apart.

Finally, we tested the requirement for fluidization at the midplane itself. Stabilizing midplane F-actin with phalloidin at 1.3 µM prevented complete F-actin disassembly without abolishing spindle dynamics (Fig. 4j–k, Extended Data Fig. 6f). This treatment slowed centrosome separation and increased the frequency of Collapse (Fig. 4l, Extended Data Fig. 6g). Myosin-II contractility thus drives aster-spanning flows specifically by contracting the gel away from the Aurora B–fluidized midplane.

### Bulk actomyosin supports long-range mechanical coupling in cytoplasm

Our emerging model depended on the ability of actomyosin to generate coherent flows over hundreds of microns, which requires long-range mechanical coupling. To test if this was physically plausible, we artificially deformed the cytoplasm and measured the length scale of responses. We inserted a 100-µm nitinol needle vertically into the extract and displaced it over 100 µm at 10 µm/s. To measure the mechanical response of cytoplasm, we tracked 1 µm PEG-coated beads (Fig. 5a; Methods). These experiments were performed during late anaphase to early interphase to ensure an organization state that matched cytoplasmic partitioning and aster separation. In microtubule-free cytoplasm with intact F-actin, deformation propagated along the path of needle displacement for hundreds of µm away from the needle in both directions, decaying with a half-length of 200–300 µm (Fig. 5b, c, Extended Data Fig. 6h). This length scale is comparable to the scale of the anaphase flows and to the radius of a *Xenopus* zygote (500–600 µm). The cytoplasm partially recoiled when the needle was lifted, revealing an elastic component (not shown). Depolymerizing F-actin with LatB destroyed long-range coupling. Needle movement produced only local, vortex-like displacement and the half-decay length shortened roughly three-fold (Fig. 5b, c, Extended Data Fig. 6h). The bulk cytoplasm therefore behaves as an F-actin-dependent crosslinked viscoelastic gel that transmits force across space, whereas the absence of F-actin converts it into a largely viscous medium in which forces dissipate locally.

**Fig. 5:**
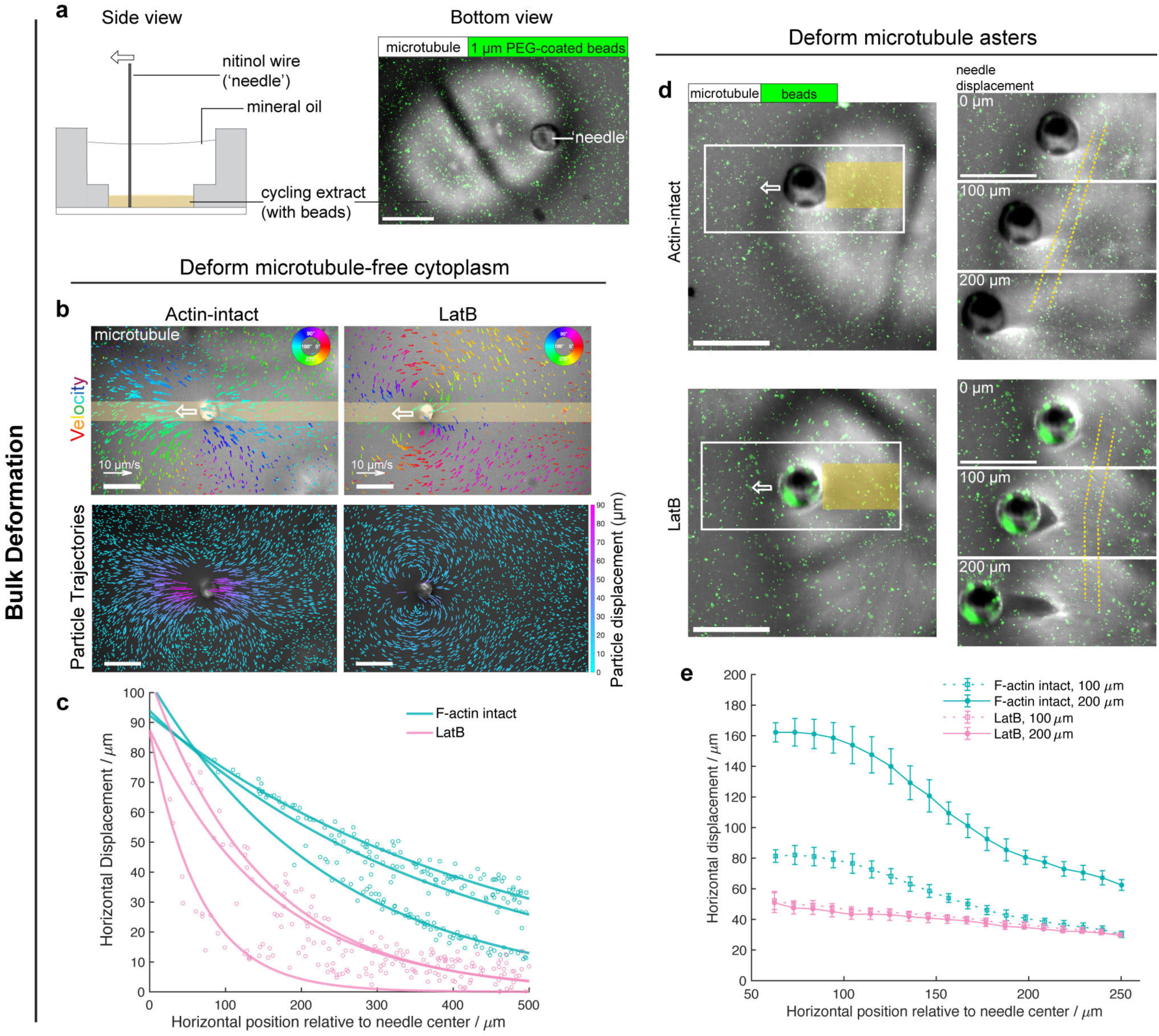
Bulk actomyosin supports long-range mechanical coupling in cytoplasm. **a**, Experimental setup of manipulating a 100 µm-thick nitinol wire (‘needle’) to deform egg extract. Left: side-view schematics. Right: bottom view showing images of microtubules (Tau-mCherry) and 1 µm PEG-coated Fluoresbrite YG beads during needle manipulation. **b**, Cytoplasmic deformation in untreated actin-intact egg extract and the same extract treated with 10 µM LatB. The needle moved 100 µm to the left in microtubule-free cytoplasm over 10 seconds during late anaphase-early interphase. Fluorescent beads were imaged at 100 ms interval and individually tracked. Top: velocity vectors of beads color-coded by orientation were overlaid on microtubule (Tau-mCherry) images captured before deformation started, showing the lack of microtubules in needle vicinity. Vectors represent velocity 5 seconds after needle movement started, averaged over 10 frames (1 second). Only 1/3 of all velocity vectors were displayed for better visualization. Bottom: particle trajectories over the entire course of the deformation, color-coded by total displacement. **c**, Horizontal displacement of beads along the path of needle movement (100 µm-wide, yellow shaded in **b**) plotted against horizontal position relative to needle center from 3 pair of experiments as in **b**. Particle displacement on the side of cytoplasm being stretched was shown and fitted to exponential decays. Also see Extended Data Fig. 6h. **d**, Microtubule asters stretched in actin-intact and 10 µM LatB-treated cytoplasm, both prepared from the same egg extract. The needle was displaced over two 100 µm steps in each experiment, at the same 10 µm/s velocity as above. Microtubules (Tau-mCherry) and beads were co-imaged at 2.6-sec interval. **e**, Quantification of aster deformation in **d** with particle image velocimetry (PIV). Horizontal displacement of beads along the path of needle movement on the side being stretched was shown (yellow shaded in **d**). PIV was performed frame-by-frame and computationally combined to obtain total displacement. Data are represented as mean ± SD. All scale bars, 200 µm.

We then asked whether microtubule asters are coupled into the actomyosin gel. Stretching an aster from its edge in two 100-µm steps, we found that with intact F-actin the aster deformed coherently as a single soft body even at 200 µm of stretch, whereas without F-actin it fractured at its periphery before 100 µm (Fig. 5d, e; Supplementary Video 8 and 9). Microtubules alone could neither hold the aster together nor transmit force across it. F-actin interwoven with microtubules supplied the mechanical strength and continuity required for smooth, long-range deformation in response to force. Bulk actomyosin thus couples asters, organelles, and cytosol into one force-propagating gel, providing the continuity required for contractile stress to move cytoplasm over hundreds of µm. Together, these data describe an active composite gel under actomyosin-mediated contractile stress with a locally fluidized midplane.

### Global contractile stress and local fluidization are sufficient to generate partitioning flows

Our data suggested a minimal physical picture where an actomyosin gel under global contractile stress is locally fluidized at the midplane and flows away from that fluidized plane (Fig. 6a). To test sufficiency of this concept over length and time scales relevant to the cytoplasmic flows, we modeled the bulk cytoplasm as an incompressible viscous continuum governed by the Stokes equation in the low-Reynolds-number regime^13^, with an active contractile stress (*s*) and a viscosity (η) each set proportional to local actomyosin density, and imposed a growing low-stress, low-viscosity zone to represent the fluidized midplane (Eqn. (1) and (2) below; Fig. 6b; Methods).

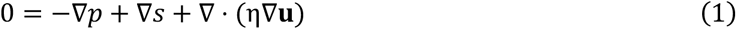

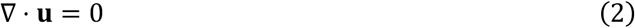

**Fig. 6:**
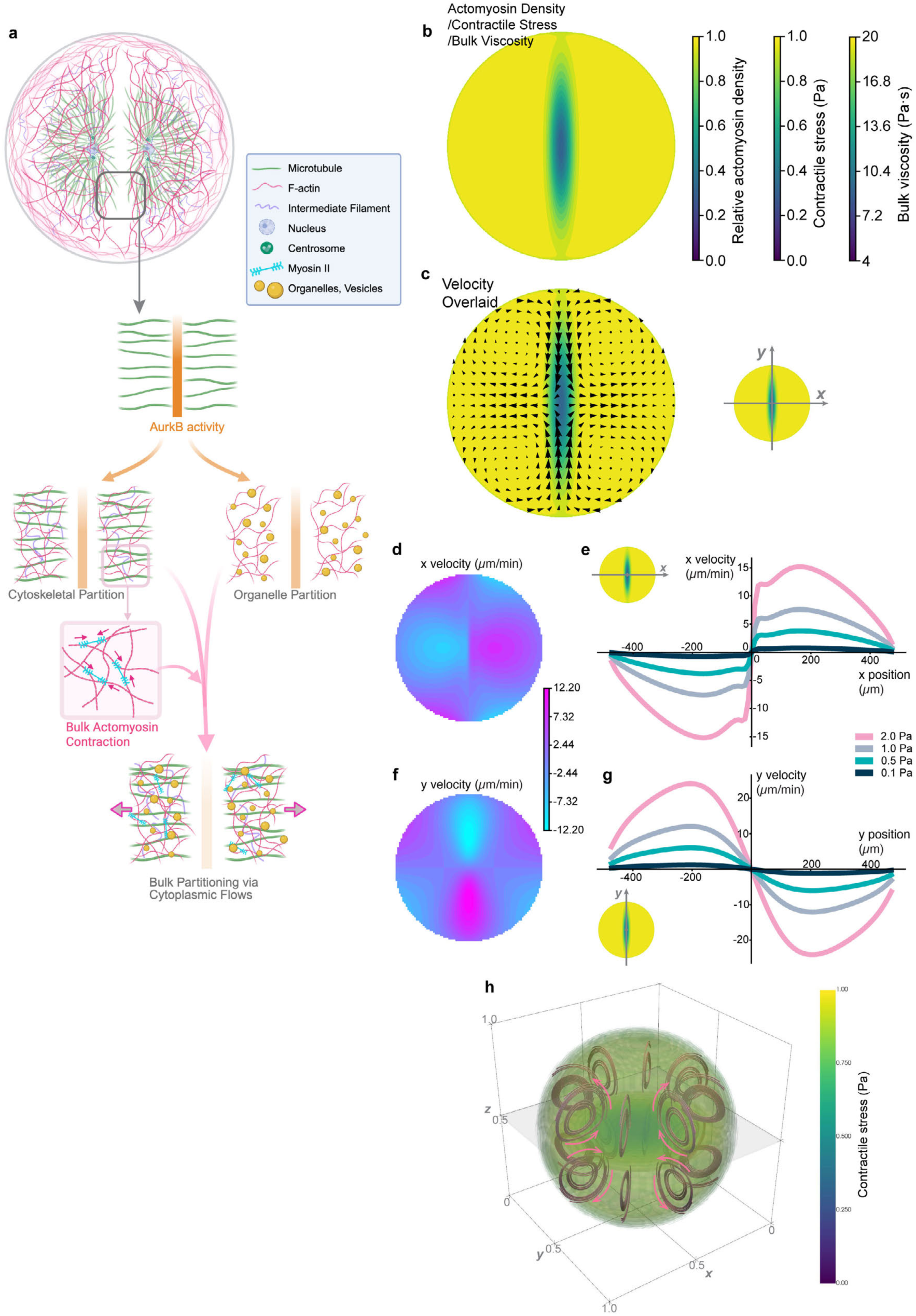
Global contractile stress and local fluidization are sufficient to generate partitioning flows. **a**, Summary of bulk cytoplasmic partitioning via actomyosin-mediated flows. Interaction between microtubule asters localizes Aurora B kinase along the midplane, whose activity locally depletes both cytoskeleton and organelles. Cytoplasmic F-actin organizes a composite gel with microtubules and organelles. Myosin-II contractility drives bulk flows that partition cytoplasm away from the fluidized midplane. **b**, Spatial setup of a 2D fluid dynamics model. A low-stress low-viscosity Gaussian ellipsoid simulated the fluidized midplane in a post-mitotic embryonic cell. Both contractile stress (0 – 1.0 Pa) and bulk viscosity (4 – 20 Pa·s) were set to be linearly dependent on local actomyosin density. Diameter of the cell was set as 1 mm. **c**, 2D simulation of cytoplasmic velocity from solving the Stokes equation with parameters shown in **b**. The *x* axis represented the separation axis, consistent with Fig. 4; the orthogonal *y* axis aligned with the midplane. **d** and **f**, Projection of velocity vectors in **c** onto the *x* and *y* axes, respectively. **e** and **g**, Cytoplasmic velocity along the *x* axis (y = 0) and along the *y* axis (x = 0), respectively, with the maximum contractile stress varying between 0.1 Pa and 2.0 Pa. **h**, 3D simulation of cytoplasmic flows with the midplane modeled as a thin disc at z = 0.5. Same parameters were used as in **b** and **c** to solve the Stokes equation in 3D.

Using bulk viscosity (4–20 Pa·s) and contractile stress (≤1–2 Pa) estimated from related embryo and extract measurements^17,18,39^, we solved Eqn. (1) and (2) with a no-slip, no-penetration boundary condition to obtain cytoplasmic velocity **u**. This model reproduced the experimental flows in both pattern and magnitude. Cytoplasm flowed away from the fluidized midplane along the separation axis (*x*) at up to ∼7 µm/min, with compensating inward flow along the midplane (*y*) forming a vortex in each quadrant (Fig. 6c–g). Flows scaled with contractile stress, persisted across the full range of physically reasonable parameters, and recurred in three-dimensional simulations (Fig. 6h, Extended Data Fig. 7). Reduced contractile stress led to slower flows, consistent with the experimental condition where myosin-II was inhibited. Imposing contractile stress with a broken symmetry on a simple fluid dynamics model is therefore sufficient to recapitulate the long-range, vortex-like flows that partition the cytoplasm.

## Discussion

Partitioning cytoplasm over large spatial scales is a fundamental physical demand of early animal development. Our results identify a mechanism for this scaling problem in which the bulk cytoplasm itself functions as an active composite gel whose fluidity is locally switchable by Aurora B activity. Actomyosin couples microtubule asters, organelles and nuclei into a mechanically cohesive, gel-like composite that transmits forces over hundreds of micrometers. After mitosis, Aurora B kinase patterns a locally fluidized midplane within this otherwise coupled cytoplasm. Myosin-II contractility then converts this local asymmetry into coherent global flows that move cytoplasmic components away from the midplane. Thus, post-mitotic partitioning is not achieved as the sum of discrete cargo-specific transport events through cytoplasm, but by cytoplasm-scale remodeling and flow of an active material.

Our model reframes a problem that has largely been approached one molecule (or organelle) at a time. Aster positioning and organelle distribution in dividing cells have mostly been explained through microtubules, dynein and cargo–motor interactions^2,6,40,41^. In embryos that are several hundred µm to 1 mm across, no individual microtubule filament reaches across the cell, and our needle-deformation and aster-stretching experiments show why a purely microtubule-based account is insufficient. It is actomyosin, not microtubules, that supplies the mechanical continuity needed to transmit force across an aster and across the bulk cytoplasm. Microtubules alone could not provide mechanical strength or propagate strain, whereas actomyosin allowed asters to behave as soft bodies embedded in a force-transmitting gel. The cytoplasm-as-material view, long developed as a physical characterization of viscoelasticity and active matter mechanics, here acquires a specific biological function: the material state of cytoplasm is the substrate on which division acts. Our data connect these two traditions by showing that a cell actively patterns its own material properties to achieve bulk partitioning.

A central feature of our proposed mechanism is that a biochemical activity sets a mechanical boundary condition in space. Aurora B kinase, as part of the chromosomal passenger complex, is known to enrich on antiparallel microtubules between neighboring asters^4^; we find that this midplane localization locally clears cytoskeleton and organelles, converting a gel-like composite into a near-viscous fluid at the midplane (α approaching 1, diffusivity roughly threefold higher). Fluidization does not itself generate directed motion, rather it defines where contractile stress is released. Symmetry breaking and force generation are thus separable. Aurora B supplies the spatial information, and bulk actomyosin supplies the mechanical power. This active stress-generating role of actomyosin is coupled to the mechanical continuity it supports, as contractile stress can generate long-range coherent cytoplasmic transport only if acting through a coupled medium. Our continuum model makes this picture explicit, reproducing the observed vortex-like flow geometry and 5-10 µm/min magnitudes from a contractile gel with a growing low-viscosity, low-stress zone. The spatial patterning role of Aurora B is consistent with the existing framework where microtubule asters dominate the geometry of cytoplasmic partitioning^21,22^, yet we argue that a mechanical perspective was missing, and actomyosin supplied physical robustness and mechanical forces on top of geometric patterning.

This division of labor between F-actin and microtubules revealed by our system may be general. Cortical actomyosin is well known to drive large-scale flows and symmetry breaking at the cell surface^42,43^, and actin-driven dynamics organize chromatin, nuclear or spindle localization in oocytes and zygotes^14–16,28,44,45^. Our work extends this logic into the three-dimensional bulk cytoplasm and shows that, as cells become very large and their surface-to-volume ratio falls, the bulk cytoplasmic gel can dominate internal mechanical organization. The same elements recurred across cycling *Xenopus* extract, *Xenopus* embryos and live medaka embryos, suggesting that coherent cytoplasmic flows away from a fluidized midplane is a conserved strategy for partitioning cytoplasm at large scale rather than a peculiarity of one system. It also offers a mechanical solution that ensures robust cleavage, as coupling all contents into one gel makes their co-segregation a material inevitability rather than a process that must be powered and regulated separately for each organelle.

## Methods

### Animal models

The *Xenopus laevis* frogs (adult females, ∼3 years old) used in these studies are part of the *Xenopus* colony bred and maintained at Harvard Medical School under the direction of Dr. Marc Kirschner. The frogs are housed in flow-through ponds with at least 1 gallon of RODI treated water per frog. The frogs receive 100% water changes 3 days a week and are allowed at least 3 months to replenish oocytes between induced egg-laying cycles. Husbandry facilities and all experimental protocols involving frogs were reviewed and approved by the Harvard Medical School’s Care and Use Committee. All studies using *Xenopus laevis* followed the guidelines of the U.S. Department of Health and Human Service for the Care and Use of Laboratory Animals, and all experiments were performed in accordance with national regulatory standards and ethical rules, Mitchison Lab IACUC Protocol number IS00003349-6.

Medaka (*Oryzias latipes*) experiments were conducted according to protocols ACUP-2023-009 and ACUP-2025-068 approved by the Animal Care and Use Committee at Okinawa Institute of Science and Technology Graduate University (OIST). The OK-Cab strain (MT830) of medaka was obtained from the National Bio-Resource Project Medaka (NBRP Medaka) and used as the parental strain. Medaka were raised and maintained as described previously^32^.

### *Xenopus* egg extract experiments

#### Preparation of actin-intact cycling *Xenopus* egg extract

Actin-intact cycling Xenopus egg extract was adapted from published cycling extract protocols^23,46^ by omitting cytochalasin during extract preparation. *Xenopus laevis* females were injected with 100 IU pregnant mare serum gonadotropin (PMSG; ProSpec, HOR-272) 3-10 days before egg laying and then with 500 IU human chorionic gonadotropin (hCG; Merck, 133754) the day before extract preparation. On the morning of extract preparation, frogs were moved to separate bins of 1x MMR (0.1 M NaCl, 2 mM KCl, 1 mM MgCl_2_, 2 mM CaCl_2_, 0.1 mM EDTA and 5 mM HEPES, pH 7.8) to lay eggs at 16 °C. Egg batches from different females were processed separately and extracts were not pooled. Fresh eggs laid within 4-5 h were collected and washed once with 0.2x MMR. All subsequent steps were performed at 18 °C with pre-cooled buffers before eggs were moved onto ice. Eggs were dejellied with two washes of 20 g/L cysteine in 1x extract buffer (100 mM KCl, 0.1 mM CaCl_2_, 1 mM MgCl_2_, 50 mM sucrose and 10 mM HEPES, pH 7.7). After washing with 0.2x MMR, eggs were activated with 0.25 µg/ml calcium ionophore A23187 (Sigma-Aldrich, C7522) in 0.2x MMR by gentle swirling in glass dishes for approximately 5 min and no longer than 7 min. Activated eggs were washed with 1x extract buffer followed by 1x extract buffer containing LPC protease inhibitors (10 µg/ml each leupeptin, pepstatin and chymostatin). Eggs were transferred to clear centrifuge tubes and packed by centrifugation at 700g for 30 s followed by 1,400g for 15 s. Eggs were then placed on ice, residual buffer was aspirated, and the eggs were crushed at 15,000g for 15 min at 4 °C. The cytoplasmic layer was collected by puncturing the tube with an 18 G needle, supplemented with 1/1,000 volume additional LPC and kept on ice. Extracts were used fresh within approximately 4 h.

#### Actin perturbations

To prepare cytochalasin D-treated cycling extract (“CytoD extract”) in parallel with actin-intact extract, two tubes were prepared from the same egg batch. 1 ml of 1x extract buffer containing LPC was loaded into each tube; 10 µl cytochalasin D stock (Cayman Chemical, 11330; 10 mg/ml in DMSO) was added to the second tube and mixed by pipetting. Activated and washed eggs were split between the two tubes, loading the actin-intact tube first to avoid cytochalasin contamination. After the crushing spin, the cytochalasin-treated extract was further supplemented with 1/1,000 volume cytochalasin D stock. Latrunculin B (Cayman Chemical, 10010631) was added to actin-intact extract at the indicated final concentrations, typically 10 µM for imaging assays and 50 µM for mitotic-outcome scoring where indicated.

#### PEG-passivated chambered coverslips and loading extract reactions

µ-Slide 15 Well 3D Glass Bottom chambered coverslips (ibidi, 81507) were passivated with PLL-g-PEG (SuSoS, PLL(20)-g[3.5]-PEG(2)) using an approach adapted from published glass passivation methods^47^. Chambers were washed with 12 µl Milli-Q water, incubated with 12 µl 2 M NaOH for 1 h, washed three times with water (12 µl, 40 µl and 40 µl) and once with 40 µl 70% ethanol. Each chamber was then incubated for 30 min with 12 µl 0.1 mg/ml PLL-g-PEG in 10 mM HEPES, pH 7.7. PLL-g-PEG was aspirated, chambers were rinsed three times with water (12 µl, 40 µl and 40 µl), and the passivated surfaces were dried by careful aspiration. Passivated chambers were stored at room temperature and typically used within 2 weeks.

Cycling extract was mixed with sperm nuclei and fluorescent probes as described below. For imaging, 1.8 µl reaction mix was placed at the center of each passivated chamber, avoiding contact between the droplet and chamber wall. The extract droplet spread slightly on the PEG-passivated surface and was then gently covered with light mineral oil to allow gas exchange and robust cell-cycle progression. This semi-open chamber allowed direct mechanical or chemical perturbation during live imaging.

#### Sperm nuclei and nuclear markers

Demembranated *Xenopus laevis* sperm nuclei were prepared from testes of male frogs as described previously^48^. Aliquots of sperm nuclei in storage buffer (30% glycerol, 0.3% BSA, 250 mM sucrose, 1 mM EDTA pH 8.0, 0.5 mM spermidine, 0.2 mM spermine, 1 mM DTT and 15 mM K-HEPES, pH 7.7) were flash frozen in liquid nitrogen and stored at -80 °C. For optimal chromatin dynamics in cycling extract reactions, sperm nuclei were used within 1 year of preparation. On the day of experiment, sperm nuclei were diluted in sperm dilution buffer (50 mM KCl, 0.5 mM MgCl_2_, 150 mM sucrose and 10 mM K-HEPES, pH 7.7) and mixed using a wide-bore pipette tip. To obtain sparsely spaced asters at the start of cycling extract reactions, sperm nuclei were used at final concentrations below 10 nuclei/µl.

GFP-NLS^49^ and NLS-2xBFP (Addgene, 176151), both expressed in *E. coli* and affinity purified, were used as nuclear markers. GFP-NLS was used at 30 nM final concentration. NLS-2xBFP was used at approximately 3.5 µM because of strong cytoplasmic autofluorescence under UV excitation, presumably deriving largely from NADH. Chromatin dynamics were visualized with JF646-Hoechst at 1 µM for DNA staining, which produced minimal perturbation from phototoxicity among a range of DNA dyes tested. JF646-Hoechst was a gift from Luke Lavis (Janelia Research Campus, Ashburn VA) to Tae Yeon Yoo (currently Harvard University, Cambridge MA).

#### Cytoskeletal and organelle markers in egg extract

Microtubules were imaged using tubulin-AF647, tubulin-CF543, Tau-mCherry or EB1-mApple in different experiments. Bovine tubulin was labelled with Alexa Fluor 647 (Thermo Fisher, A20106) or CF 543 (Biotium, 92105) as described previously^4,50^ and used at 50 nM. Tau(mTMBD)-mCherry was expressed and purified as described previously^51^ and used at 200-400 nM. EB1-mApple^52^, which binds microtubule plus ends and improves visualization of centrosomes relative to labelled tubulin or Tau-mCherry, was used at 200 nM. F-actin was imaged with Lifeact-GFP at 500 nM to 1 µM^34^. Keratin was imaged with a mouse monoclonal anti-pan cytokeratin (Sigma-Aldrich, P2871) antibody directly labelled with CF647 (Biotium, 92135) and used at 4 µg/ml. CPC localization was imaged with directly labelled anti-INCENP where indicated^34^.

Acidic organelles, including lysosomes and endosomes, were labelled with LysoTracker Deep Red (Invitrogen, L12492) at 300-500 nM. Mitochondria were labelled with MitoView 405 (Biotium, 70070) at 530 ng/ml, MitoView Fix 640 (Biotium, 70082) at 100 ng/ml or DiOC6(3) (Sigma-Aldrich, 318426) at 600 ng/ml (1 µM), depending on the experiment. ER was labelled with Vybrant DiD (Invitrogen, V22889) at 400 nM using a pre-incubation step to improve uniform labelling. Vybrant DiD stock (1 mM in ethanol-containing solvent) was diluted 1:50 into fresh egg extract and incubated at 18 °C for more than 1 h, and the pre-incubated DiD-extract mixture was diluted 50-fold into the final extract reaction.

#### Direct antibody labelling

IgG labelling with fluorescent dyes was performed either on resin using Protein G UltraLink resin (Thermo Scientific, 53125) or Affi-Prep Protein A resin (Bio-Rad, 156-0006), as described previously^47^, or in solution. Fluorescent dyes we used include Alexa Fluor 488, 568 and 647 NHS esters (Thermo Fisher Scientific, A20100, A20003 and A20106) and CF543 and CF647 NHS esters (Biotium, 92105 and 92135). For in-solution labelling, anti-keratin antibody was mixed on ice with 1/10 volume 1 M K-HEPES, pH 7.7, and 1/40 volume CF647 NHS ester (10 mM in DMSO). The reaction was incubated at room temperature for approximately 2 h and gel filtered using a NAP-5 column containing Sephadex G-25 (Cytiva, 17085301) equilibrated in 100 mM KCl, 10 mM K-HEPES, pH 7.7, and 1 mM EGTA at 4 °C. Fractions containing labelled antibody were identified by thin-layer chromatography and ultraviolet-visible absorbance using a NanoDrop spectrophotometer.

#### PEG-passivated fluorescent beads

Fluoresbrite 641 carboxylate microspheres (0.5 µm; Polysciences, 21116-1) and Fluoresbrite YG carboxylate microspheres (0.5 or 1.0 µm; Polysciences, 21636-1), supplied as 2.5-2.7% aqueous suspensions, were covalently coated with PEG by EDC/Sulfo-NHS coupling. Beads were pelleted and washed twice with 0.2 M sodium phosphate buffer, pH 7.2, and then resuspended with α-methoxy-ω-amino PEG (CH3O-PEG-NH2, 2,000 Da; Rapp Polymere, 122000-2) at 100 mg/ml (50 mM) in sodium phosphate buffer. Freshly prepared N-hydroxysulfosuccinimide sodium salt (Sulfo-NHS; Sigma-Aldrich, 56485) and N-(3-dimethylaminopropyl)-N′-ethylcarbodiimide hydrochloride (EDC; Sigma-Aldrich, E7750) were added to final concentrations of 5 mM and 100 mM, respectively. Suspensions were mixed thoroughly and rotated overnight at room temperature. Beads were then pelleted, washed and stored in 100 mM KCl and 10 mM K-HEPES, pH 7.7, at 4 °C. After passivation, stocks were diluted more than ten-fold and contained approximately 10^9^ – 10^10^ particles/ml. For flow tracking, microrheology and needle-deformation assays, PEG-passivated beads were diluted approximately 1:100 into extract, giving final concentrations of approximately 10^7^ – 10^8^ particles/ml.

#### Needle deformation of egg extract

Super-elastic nitinol wire of 0.004-inch diameter (approximately 100 µm; McMaster-Carr, 8320K11) was used as a needle to deform extract. A custom micromanipulator was assembled from two linear stages with a 0.635-mm leadscrew pitch (Applied Scientific Instrumentation, LS-25-UMCRX), coupled vertically using a dovetail mount and controlled with a multi-axis stage controller (Applied Scientific Instrumentation, MS2). The assembly was mounted on a vibration-isolation air table using Thorlabs components. A steel threaded rod mounted on the linear stages was connected to a short segment of nitinol wire through a sterile 30 G needle (BD, 305128). Python scripts controlled the stages to impose specified displacements and velocities. For the needle-deformation experiments in Fig. 5, the needle was inserted vertically into extract and displaced by 100 µm at 10 µm/s. PEG-passivated Fluoresbrite YG beads were imaged alone at 10 Hz for particle tracking, or together with Tau-mCherry at 2.6-s intervals when aster deformation was measured.

#### Inhibitory antibody against MYH9

A C-terminal fragment of *Xenopus* MYH9 (aa 1720-1925), corresponding to the assembly-critical domain (ACD) required for myosin-II mini-filament assembly, was cloned into pET-28a to generate a C-terminal His-tagged construct (Twist Bioscience). The plasmid was propagated in NEB Turbo competent E. coli (New England Biolabs, C2984H) and purified using a QIAprep Spin Miniprep Kit (Qiagen, 27104). The His-tagged construct was cut with a restriction enzyme and used to assemble a C-terminal GST-tagged construct with Gibson Assembly Master Mix (New England Biolabs, E2611S). MYH9_ACD_-His and MYH9_ACD_-GST were expressed in BL21(DE3) competent E. coli (New England Biolabs, C2527I) and purified using Ni-NTA agarose (Qiagen, 30230) and Pierce Glutathione Agarose (Thermo Scientific, 16100), respectively.

The MYH9 assembly-critical domain (MYH9_ACD_) sequence used for immunization was: ALEEKRRLESRISQLEEELEEEQGNTELVNDRLKKATLQIDQMNADLNAERSNAQKNENARQQMDRQNKELKTKLQEMEGTIKSKFKANITALEAKIAQLEEQLDSETKERQNASKQVRRTEKKLKDVMIQVEDERRNSEQYKDQAEKNNVRMKQLKRQVEEAEEEAQRANAMRRKLQRELEDATETAEAMNREVNTLKTKLRRGG

MYH9_ACD_-GST was used as antigen to immunize a rabbit (Cocalico Biologicals). MYH9_ACD_-His was coupled covalently to Affi-Gel 10 (Bio-Rad, 1536099) through NHS chemistry and used as the affinity column. Rabbit serum containing anti-MYH9_ACD_-GST was incubated overnight at 4 °C with the MYH9_ACD_-His column equilibrated in 1× TBS. The resin was washed sequentially with 1× TBS, 5× TBS and 1× TBS, and anti-MYH9_ACD_ was eluted with 0.2 M acetic acid and 0.2 M NaCl (pH ∼2.6). The purified antibody was buffer exchanged using Amicon Ultra 100-kDa filters into 100 mM KCl and 10 mM K-HEPES, pH 7.7, and concentrated to 42 mg/ml. ChromPure whole-molecule rabbit IgG (Jackson ImmunoResearch, 011-000-003) was exchanged into the same buffer and concentration as the non-specific control. Antibodies were used at 2 mg/ml final concentration by adding 0.5 µl antibody to 10 µl egg extract containing imaging probes and sperm nuclei.

To validate anti-MYH9 specificity, egg extract was diluted 20-fold in LDS sample buffer, denatured at 80 °C for 10 min, separated by gel electrophoresis and transferred to a cellulose membrane. Western blotting was performed using purified anti-MYH9_ACD_. To test inhibition of myosin-II-dependent bulk contraction, Lifeact-GFP was added to cycling egg extract and non-specific rabbit IgG or anti-MYH9ACD was added to separate aliquots at 2 mg/ml final concentration. Light mineral oil (Sigma-Aldrich, 330779; 10 µl) was loaded into unpassivated glass-bottom chambers (ibidi, 81507), and 1 µl extract reaction was pipetted into the oil and allowed to settle to the bottom because of its higher density. Extract droplets were imaged with a 4× objective for 2 h to assay bulk contraction.

#### F-actin stabilization with phalloidin

Phalloidin (Cayman Chemical, 18039) stock (10 mg/ml in DMSO, 12.7 mM) was diluted 20-fold into actin-intact egg extract to 0.5 mg/ml and frozen in aliquots. On the day of the experiment, the frozen phalloidin-in-extract stock was thawed and diluted into fresh extract. To reach 2.5 µM final concentration, the 0.5 mg/ml stock was diluted 50-fold with fresh extract to 10 µg/ml and then diluted a further 5-fold to 2 µg/ml (2.54 µM). To reach 1.3 µM final concentration, the 0.5 mg/ml stock was diluted 10-fold with fresh extract to 50 µg/ml and then diluted a further 50-fold to 1 µg/ml (1.27 µM). The morphology of cytoplasmic F-actin under phalloidin treatment varied with the dilution scheme, especially at intermediate concentrations where F-actin was partially stabilized.

#### Aurora B inhibition with barasertib-HQPA

Barasertib-HQPA (MedChemExpress, HY-10126) was prepared as an aqueous stock rather than a DMSO stock to avoid perturbing microtubule dynamics by DMSO addition. A 10 mM aqueous stock was prepared by dissolving barasertib-HQPA in 18 mM acetic acid. For partial inhibition at 0.5 µM final concentration, a small volume of aqueous stock was rapidly mixed with egg extract to make a 1:50 dilution, avoiding precipitation during the pH change, and then diluted a further 400-fold into extract. For timed local addition, barasertib-HQPA was rapidly diluted 20-fold with egg extract to 0.5 mM. Untreated cycling extract was allowed to progress to anaphase, and 0.2-0.3 µl of 0.5 mM barasertib-HQPA was pipetted into the field of view adjacent to aster pairs of interest.

#### Egg extract microscopy

Unless otherwise stated, egg extract imaging was performed by widefield microscopy on a Nikon Eclipse Ti2-E inverted microscope equipped with a Lumencor SOLA SE V-nIR light engine and an Andor Zyla 4.2 PLUS sCMOS camera. Images were acquired as z-stack time lapses using either a Nikon CFI Plan Apo Lambda 10×, NA 0.45 objective or a Plan Apo Lambda 20×, NA 0.75 objective. Four fluorescence channels were used: DAPI (Nikon, 96359; excitation 350/50, emission 460/50), GFP (Nikon, 96362; Chroma, 49002 ET GFP; excitation 470/40, emission 525/50), Texas Red (Nikon, 96365; excitation 560/40, emission 630/75) and Cy5 (Chroma, 49009 ET Cy5 NX; excitation 640/30, emission 690/50). Images were acquired using NIS-Elements software (Nikon). The microscope room was maintained at 18 °C. Confocal imaging of the extract experiment shown in Fig. 1d was performed at the Harvard Medical School Core for Imaging Technology & Education (CITE) using a Nikon Ti2-E motorized inverted microscope with Perfect Focus, a Nikon AX R scanning head with Galvo and Resonant scanners, and NIS-Elements software. Images were acquired using a Plan Fluor 20× MImm DIC N2, NA 0.75 objective and a Plan Apo Lambda 60× oil, NA 1.4 objective.

### *Xenopus* embryo experiments

#### *Xenopus* embryo fertilization and drug treatment

*Xenopus laevis* females were induced to ovulate with PMSG (ProSpec, HOR-272) and hCG (Merck, 133754) as described for extract preparation. Testes were isolated from male frogs as described previously^48^ and used the same day or stored at 4 °C for up to 1 week. Eggs were gently squeezed from laying females into a dry Petri dish and fertilized immediately by passing a small piece of macerated testis over all eggs. After 5 min incubation, the Petri dish was flooded with 0.1× MMR. All steps were performed at 18 °C.

For latrunculin B treatment and fixation, zygotes were dejellied at approximately 15 min post-fertilization by adding 2% cysteine in 0.1× MMR and swirling for 5-7 min. They were then rinsed five times with 0.1× MMR and transferred at 20 min post-fertilization to Petri dishes containing either 0.3× MMR as control or 10 µg/ml latrunculin B (Cayman Chemical, 10010631) in 0.3× MMR. After gentle swirling for 1 min, embryos were incubated in the respective buffer at 18 °C. Developing embryos were removed from buffer and fixed at 5-10-min intervals, beginning at 70-80 min post-fertilization, in methanol/EGTA fixative (50 mM EGTA, pH 6.8, 10% water and 90% methanol) with gentle agitation for approximately 24 h at room temperature. After fixation, embryos were stored in fixative at 4 °C. In some experiments, except those involving CPC imaging, embryos were pre-fixed for 5 min at 18 °C with gentle shaking in PFA/glutaraldehyde fixative (0.50% paraformaldehyde prepared from 16% stock (Ted Pella, 18505), 0.1% glutaraldehyde, 80 mM PIPES, pH 6.8, 1 mM MgCl2, 10 mM EGTA and 0.2% Triton X-100), followed by overnight methanol/EGTA fixation.

#### *Xenopus* embryo immunofluorescence, clearing and imaging

Fixed embryos were rehydrated through 25/75%, 50/50%, 75/25% and 100/0% TBS/methanol, with each step lasting 30-45 min under gentle rotation. TBS contained 50 mM Tris, pH 7.6, and 150 mM NaCl. Rehydrated zygotes were bleached for 5 h to overnight in freshly prepared bleaching solution containing 1% hydrogen peroxide (Sigma-Aldrich, 216763), 5% formamide (Sigma-Aldrich, F9037) and 0.5× SSC (75 mM NaCl and 8 mM sodium citrate, pH 7). Embryos were rinsed three times in TBS and blocked in TBSN (10 mM Tris-Cl, pH 7.4, 155 mM NaCl, 1% IGEPAL CA-630 (Sigma-Aldrich, I8896), 1% BSA, 2% fetal calf serum and 0.1% sodium azide) for at least 1 h. These steps were performed at room temperature. Embryos were then incubated for at least 24 h at 4 °C with very gentle rotation in directly labelled antibodies diluted in TBSN to 1 µg/ml. After antibody incubation, embryos were washed in TBSN at 4 °C for approximately 24 h with several solution changes.

ER was labelled with a rabbit polyclonal antibody against a cytosolic fragment of *Xenopus* lunapark (cytLnp; aa 99-441; gift from Tom Rapoport^53^), directly labelled with Alexa Fluor 568 NHS ester (Thermo Fisher Scientific, A20003). Microtubules were labelled with mouse monoclonal anti-α-tubulin clone B-5-1-2 (Sigma-Aldrich, T6074), directly labelled with Alexa Fluor 568 or 647 NHS ester (Thermo Fisher Scientific, A20003 and A20106). Keratin was labelled using the same CF 647-labelled anti-pan-cytokeratin antibody (Sigma-Aldrich, P2871) used in extract experiments. The CPC was labelled using the same Alexa Fluor 647-labelled anti-INCENP antibody described above.

For clearing, embryos were washed three times in phosphate-buffered saline and dehydrated through 30/70%, 50/50%, 70/30% and 100/0% 1-propanol/PBS using 1-propanol (Sigma-Aldrich, 279544), with each step lasting 30-45 min at room temperature with gentle agitation. Approximately 1.5 ml ethyl cinnamate (Sigma-Aldrich, W243000) was loaded into a 2-ml tube, and embryos in a small volume of 1-propanol were placed gently on the ethyl-cinnamate surface without disturbing the interface. As embryos cleared, they sank to the bottom and became transparent. The 1-propanol was removed from the top of the tube. Cleared embryos in ethyl cinnamate were mounted in 1.2-mm-thick metal slides with a central hole; a coverslip was attached to the bottom with heated parafilm and another coverslip was placed on top. Fixed-embryo imaging was performed at the Harvard Medical School Core for Imaging Technology & Education using a Nikon Ti-E inverted microscope with a Nikon A1R point-scanning confocal head and a 10× dry objective. Images were acquired as z-stacks through the embryo.

### Medaka fish assays

#### Exogenous protein expression in medaka embryos

To express exogenous proteins, plasmids containing Lifeact-EGFP (pTK1027), mCherry-Ol-Aurora B (pTK1072), and BiP-mCherry-KDEL (an ER marker, pTK1113)^7^ were created using a pCS2+ vector for mRNA synthesis. These plasmids were linearized by NotI or BssHII digestion, followed by *in vitro* transcription using an mMessage mMachine SP6 kit (Invitrogen, AM1340) according to the manufacturer’s instructions. Synthesized RNAs were purified with a RNeasy Mini kit (Qiagen).

#### Microinjection and live imaging in medaka embryos

HiLyte 647-tubulin (Cytoskeleton, Inc. TL670M) and mRNAs were mixed with final concentrations of 800 ng/μL for tubulin and150 ng/µL for each mRNA. The mixture was injected into one-cell embryos using siliconized glass-needles. To track cytoplasmic flows, PEG-passivated Fluoresbrite 641 beads of 0.5 μm diameter (as described above for egg extract; further 1:50 diluted with water) were injected in one of 2-cell blastomeres of EGFP-α-tubulin expressing embryo^32^. Microtubules and beads were excited by 488 nm and 640 nm laser, respectively, and visualized using a spinning-disc confocal microscope with a 20× 0.95 numerical aperture objective lens (APO LWD 20× WI λS, Nikon, Tokyo, Japan). A CSU-W1 confocal unit (Yokogawa Electric Corporation, Tokyo, Japan) with three lasers (488, 561, 640 nm, Coherent, Santa Clara, CA) and an ORCA-Fusion digital CMOS camera (Hamamatsu Photonics, Hamamatsu City, Japan) were attached to an ECLIPSE Ti2-E inverted microscope (Nikon) with a perfect focus and a water-supply system. Images were captured using NIS-Elements software (version 5.21.00, Nikon). Samples were imaged at room temperature (24-25°C).

### Data processing and image analysis

#### Mitotic outcome scoring in cycling extract

Mitotic outcomes were scored from nuclear and centrosome dynamics in cycling egg extract. To minimize variability from starting conditions, high nuclear-centrosome density and global flows, only cell-free divisions occurring at the second and third mitoses after loading into chambers were scored, and only events with no closely adjacent nuclei or centrosomes and no strong global flows were included. Scored events started with one nucleus and two attached centrosomes at prophase and were assigned to four categories: Success, Error, Collapse and Failure. Success was defined as formation of two daughter nuclei accompanied by two pairs of centrosomes that remained separated until two spindles formed in the next cycle. Failure was defined as no visible nuclear segregation or centrosome separation into daughter compartments. Collapse was defined as initial separation of both daughter nuclei and centrosomes followed by collapse before the next mitotic onset, forming a multipolar spindle. All other outcomes that did not produce a pair of daughter nuclei each attached to two centrosomes were scored as Error, including abnormal centrosome numbers, chromatin segregation failure and nuclear-centrosome detachment.

#### Quantification of organelle density

Organelle depletion and partitioning from the midplane were quantified from fluorescence intensity profiles across pairs of daughter nuclei-centrosomes during late anaphase to early interphase. For aster pairs in egg extract, 50-µm-wide lines spanning both asters were drawn in Fiji/ImageJ^54^, and intensity profiles averaged across the line width were generated using Analyze > Plot Profile. To compare channels, intensity profiles were processed in MATLAB (MathWorks) by normalizing to background intensity within asters, averaged from the 0-25% and 75-100% regions along the profile. To quantify temporal dynamics in actin-perturbed extract, 50-µm-wide and 600-µm-long lines across centrosomes were used to generate intensity profiles over time and were normalized similarly. For medaka and *Xenopus* embryos, intensity was quantified between daughter nuclei or centrosomes and normalized similarly, except that a 20-µm-wide line was used for medaka because of the smaller blastodisc geometry.

#### Cytoplasmic flow analysis

For flow analysis in egg extract, 0.5-µm PEG-coated Fluoresbrite 641 beads (Polysciences, 21116-1) and EB1-mApple were co-imaged from anaphase to early interphase at 10-s intervals using three z-planes with 5-µm spacing. Maximum intensity projections were generated for both channels. TrackMate^55^ in Fiji/ImageJ was used to track individual beads after filtering by total intensity to remove aggregates, and only trajectories lasting longer than 1 min were retained. Centrosomes were tracked semi-automatically with p53Cinema^56^ by setting the cell-size parameter to the approximate centrosome size. For each frame N, mean velocity of individual beads and centrosomes was calculated in MATLAB (MathWorks) from displacement between frames N−2 and N+2, corresponding to 40 s. The axis connecting daughter centrosomes was assigned as the *x* axis, with the origin at the midpoint between centrosomes or centrosome pairs. The orthogonal axis aligned with the midplane was assigned as the *y* axis. Velocities of centrosomes and beads within a 100-µm-wide band along the *x* axis were projected onto the *x* axis and plotted against *x* position. Local regression was performed with LOWESS in MATLAB. Similarly, velocities of beads within a 50-µm-wide band along the midplane were projected onto the *y* axis. For ensemble profiles of x velocity, asters of similar radius (270-330 µm) were normalized to 300 µm for aggregation; bead velocity was normalized so that bulk movement was set to zero. Local regression curves from individual experiments were pooled and averaged.

For flow measurements in medaka embryos, maximum intensity projections were generated from 11 z planes with 2 µm spacing, and beads were tracked with TrackMate. Because of the lower signal-to-noise ratio for microtubules imaged with EGFP-tubulin in live medaka, centrosomes were first roughly tracked using p53Cinema and then localized by two-dimensional Gaussian fitting. Subsequent flow analyses were performed as described for egg extract.

#### Pairwise MSD microrheology

For particle-tracking microrheology, 0.5-µm PEG-coated Fluoresbrite 641 beads (Polysciences, 21116-1) were imaged at 100 ms intervals for 2 min. TrackMate in Fiji/ImageJ was used to track individual beads after filtering by total intensity to remove aggregates, and only trajectories lasting longer than 4 s were retained. Pairwise mean-squared displacement (MSD_pair_) was defined as < |***r*_1_** − ***r*_2_**|^2^ >, where ***r*_1_** and ***r*_2_** are the displacements of two beads over a time lag *τ*. To compare diffusion behaviors among the 3 cytoplasmic regions shown in Fig. 3e and f, MSD_pair_ was calculated for all pairs of temporally overlapping trajectories within each region. Ensemble averages of MSD_pair_(*τ*) were plotted with SEM (invisible due to their extremely small magnitude). For regional comparisons in Fig. 3g, small areas approximately 65 × 65 µm^2^ were sampled from each cytoplasmic category, and temporally overlapping trajectories within each small area were used to calculate MSD_pair_. Ensemble averages of MSD_pair_(*τ*) within each small area were fit with log(MSD_pair_) = log(8D) + αlog(*τ*) to estimate apparent diffusion coefficient D and anomalous exponent α.

#### Analysis of needle-induced deformation

TrackMate in Fiji/ImageJ was used to obtain individual bead trajectories from 10 Hz movies acquired during needle movement. Only bead trajectories covering the entire duration of needle movement were included. Velocity vectors during deformation shown in Fig. 5b top panels represent particle velocity 5 s after needle movement began, averaged over 10 frames (1 s). Bead trajectories within a 100-µm-wide band along the path of needle movement were further analyzed by plotting horizontal displacement against horizontal position. Displacement-distance scatter plots were fit with exponential decay for visualization, based on theoretical results for related gel-deformation setups^57^. For experiments in which microtubules and beads were co-imaged at longer intervals, particle tracking failed because of the lower temporal resolution. PIVlab^58^ in MATLAB was therefore used to map deformation frame by frame, and deformation fields from successive image pairs were computationally combined to obtain total displacement across the field of view.

#### Quantification of centrosome separation

To avoid variability caused by interactions with neighboring asters, only isolated aster pairs derived from isolated spindles and lacking closely adjacent asters, especially along the centrosome separation axis (*x* axis), were analyzed for centrosome separation. Centrosomes were tracked with p53Cinema as described above. Distance between centrosomes, or later centrosome pairs, was calculated over time. To align timing of centrosome separation between experiments, time 0 was set to the first frame in which spindles were assembled and nuclei were fully disassembled.

### Fluid dynamics model

The embryonic cytoplasm was modelled as an incompressible Stokes fluid with an additional contractile stress term *s* generated by actomyosin, taking the same form as hydrostatic stress. At each position and time t, velocity **u** was modelled as:

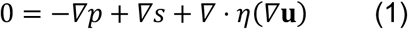

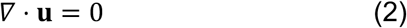

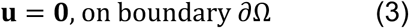

Here *η* is dynamic viscosity and *p* is dynamic pressure arising from fluid movement, which was solved rather than specified. The incompressibility condition (2) enforces conservation of mass, and the boundary condition (3) sets velocity to zero at the cell boundary.

Contractile stress *s* and viscosity *η* were each modelled as functions of normalized F-actin concentration *c*_actin_. Specifically, *s*(***x***, *t*) = *s*_0_*c*_actin_(***x***, *t*) and *η*(***x***, *t*) = *η*_max_*c*_actin_(***x***, *t*) + *η*_min_(1− *c*_actin_(***x***, *t*)). The actin depletion zone observed experimentally was represented by a growing Gaussian ellipsoid: 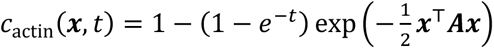, where ***A*** determines the shape and orientation of the depleted zone and (1 – *e*^−*t*^) controls growth. At t = 0, F-actin concentration is uniform; at large t, concentration contains an ellipsoidal gap centered at the midplane. This form was chosen for simplicity; the qualitative behavior depended on the existence of a gap in *c*_actin_ with a spatial gradient rather than on the specific parameterization. A spherical boundary 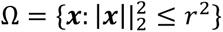 was used to represent the embryo boundary. The Stokes equation was solved numerically with the finite-volume method implemented in FiPy^59^.

Two-dimensional and three-dimensional simulations used maximum dynamic viscosity *η*_max_ = 20 Pa·s (range tested, 10-40 Pa·s)^11,17,18^, minimum dynamic viscosity *η*_min_ = 4 Pa·s (range tested, 3-8 Pa·s)^14,15,55^, maximum contractile stress *s*_0_ = 1 Pa (range tested, 0.5-2 Pa)^39^, and cell radius r = 0.5 mm. The two-dimensional matrix ***A*** that set the shape of the actin depletion zone was [[25, 0], [0, 900]], producing an aspect ratio of 6:1. In three dimensions, the depletion zone was represented as a disc along the x-y plane with aspect ratio x:y:z = 6:6:1. The stress parameter *s*_0_ was varied to simulate myosin-II inhibition. Simulations and figure generation used custom Python code, available at https://github.com/YunyiShen/Actomyosin-Flows.

## Acknowledgements

This work was supported by funding from the National Institute of General Medical Sciences (R35 GM131753 to T.J.M.) and JST FOREST (JPMJFR224O to T.K.). We thank the Mitchison group for their input and comments and Kathy Buhl and Taylor Rebbe for managing the lab. We thank Nancy Kleckner, Radhika Subramanian, Andrew Murray, L. Mahadevan, Ming Guo, James Pelletier and Rui Fang for helpful discussions. We are grateful to James Pelletier, Tae Yeon Yoo and Keisuke Ishihara for their contribution to some of the purified proteins and reagents used in this study. We thank Margaret Coughlin for the EM images. We thank the Core for Imaging Technology & Education (CITE) at Harvard Medical School (HMS) for assisting with confocal imaging, Rachael Jonas-Closs for excellent *Xenopus* husbandry, and Ofer Mazor of HMS Instrumentation Core for helping set up the micromanipulator. Elements in the schematics in Figures 1A, 5A, 5F, 6A and 7A-B were created with https://BioRender.com with licenses to L.B. The authors acknowledge the MIT Office of Research Computing and Data for providing computing resources that have contributed to the research results reported within this paper.

## Author Contributions

Conceptualization, L.B., T.J.M., C.M.F. and T.K.; Investigation, L.B., C.M.F. and A.K.; Data curation and Visualization, L.B. and T.K.; Formal analysis, L.B. and Y.S.; Methodology, L.B., C.M.F., T.J.M., A.K., T.K., Y.S. and N.O.; Resources: L.B., C.M.F., A.K., T.K. and T.J.M.; Software: L.B., Y.S. and N.O.; Supervision: T.J.M. and T.K.; Writing – original draft, L.B.; Writing – review & editing, L.B., T.J.M., C.M.F., T.K., A.K., Y.S. and N.O.; Funding acquisition, T.J.M. and T.K.

## Declaration of interests

The authors declare no competing interests.

## Extended Data

Extended Data Fig. S1 – S7

Supplementary Videos S1 – S10

## Extended Data

**Extended Data Fig. 1:**
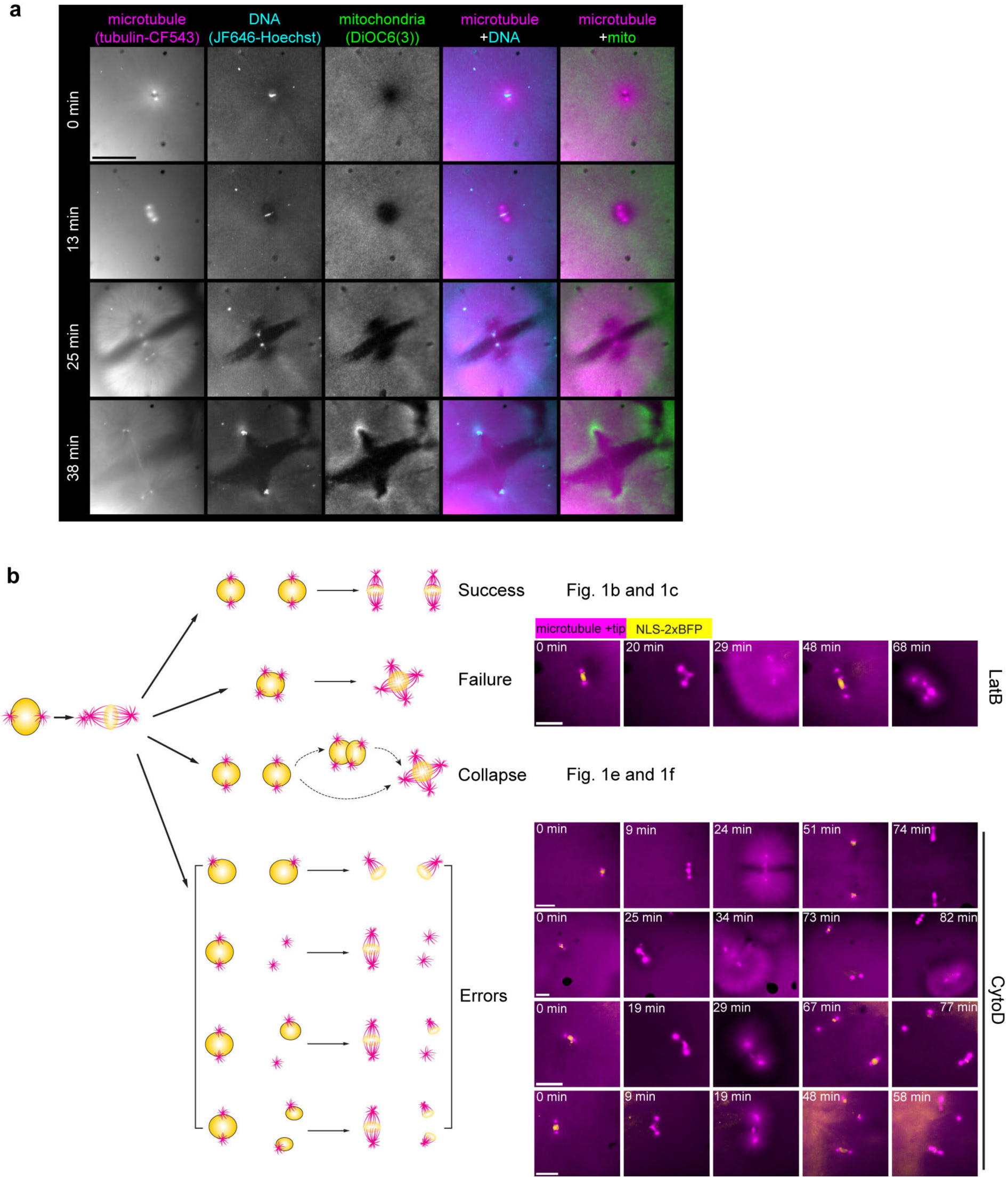
Cell cycle dynamics and mitotic errors in cycling egg extract, related to Fig. 1. **a**, Live images of DNA and mitochondria in actin-intact cycling egg extract, showing DNA segregation and mitochondria partitioning. Scale bar, 200 µm. **b**, Categories of mitotic outcomes in cycling egg extract. Microtubules and centrosomes were visualized with EB1-mApple, which binds to microtubule plus ends. Nuclei visualized with NLS-2xBFP. Scale bars, 100 µm.

**Extended Data Fig. 2:**
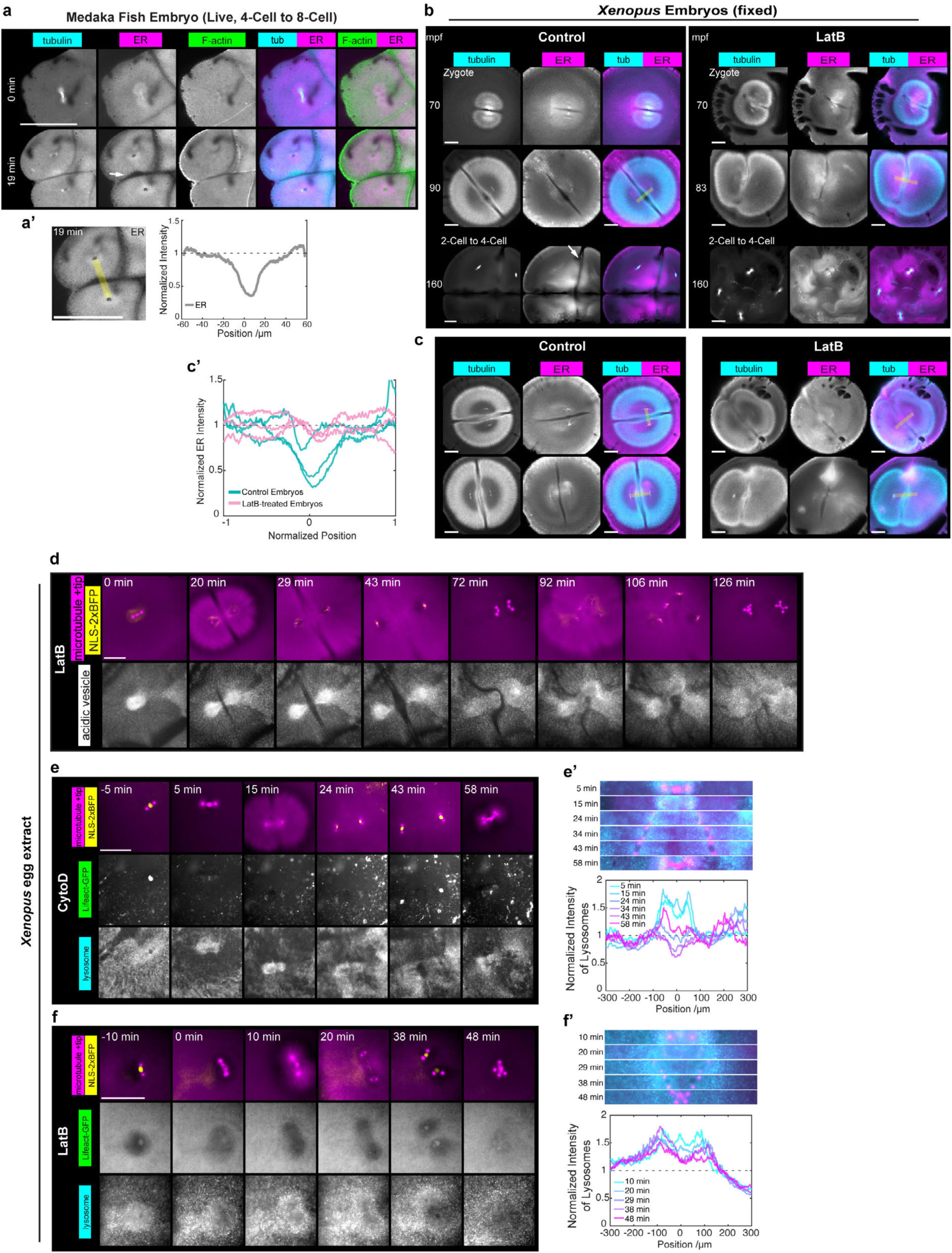
Organelle partitioning in embryos and egg extract, related to Fig. 2. **a**, Live images showing microtubules (HiLyte647-tubulin, injected), ER (BiP-mCh-KDEL) and F-actin (Lifeact-EGFP) during the 4-cell-to-8-cell mitosis in a medaka fish embryo. Arrow: ER depletion at the cleavage plane. **a’**, ER density quantified for **a**, averaged over a 20-µm-wide band between daughter nuclei (yellow shaded). **b**, Immunofluorescence images of fixed *Xenopus laevis* embryos viewed from the animal pole, visualizing microtubules (anti-tubulin-AF647) and ER (anti-cytLnp-AF568). Left: untreated embryos fixed at 70-, 90-, and 160-minutes post-fertilization (mpf). Images at 160 mpf are maximum intensity projection (MaxIP) from 3 Z-stacks. Right: embryos treated with 10 µg/ml (25 µM) LatB from 20 mpf, then fixed at 70, 83, and 160 mpf. Images at 160 mpf are MaxIP from 4 Z-stacks. Embryos fixed at 160 mpf are re-used from Fig. 2g for showing the time course development. **c**, More examples of *Xenopus laevis* embryos fixed at similar stages (80-90 mpf) as the second row of **b**, visualizing microtubules (anti-tubulin-647) and ER (anti-LNPK-AF568). Left: untreated embryos. Right: embryos treated with 10 µg/ml (25 µM) LatB. **c’**, ER density quantified for *Xenopus* embryos fixed at 80-90 mpf, untreated and LatB-treated, including one pair from the second row of **b** and two pairs from **c**. Fluorescence intensity was averaged over a 50-µm-wide band between centrosome pairs (yellow shaded in **b**). **d**, Live imaging over 2 hours of microtubules (EB1-mApple), nuclei (NLS-2xBFP) and acidic vesicles (Lysotracker Deep Red) in cycling egg extract treated with 10 μM LatB. Note the aggregation of acidic vesicles around centrosomes and variable partitioning behaviors, including unstable or poor partitioning after mitosis. **e** and **f**, Live images of microtubules (EB1-mApple), nuclei (NLS-2xBFP), F-actin (Lifeact-GFP) and acidic vesicles (Lysotracker Deep Red) in cycling egg extract prepared with CytoD (**e**) or treated with 10 µM LatB (**f**), showing collapse after initial separation. The absence of F-actin under LatB treatment is seen from diffuse Lifeact-GFP signal. **e’** and **f’**, Temporal dynamics of centrosomes and acidic vesicles across one cell cycle (top) and quantification of vesicle intensity across centrosomes over time (bottom) for the experiments in **e** and **f**. All scale bars, 200 μm. For **d**-**f**, time 0 indicates the first frame with assembled spindles (and fully disassembled nuclei).

**Extended Data Fig. 3:**
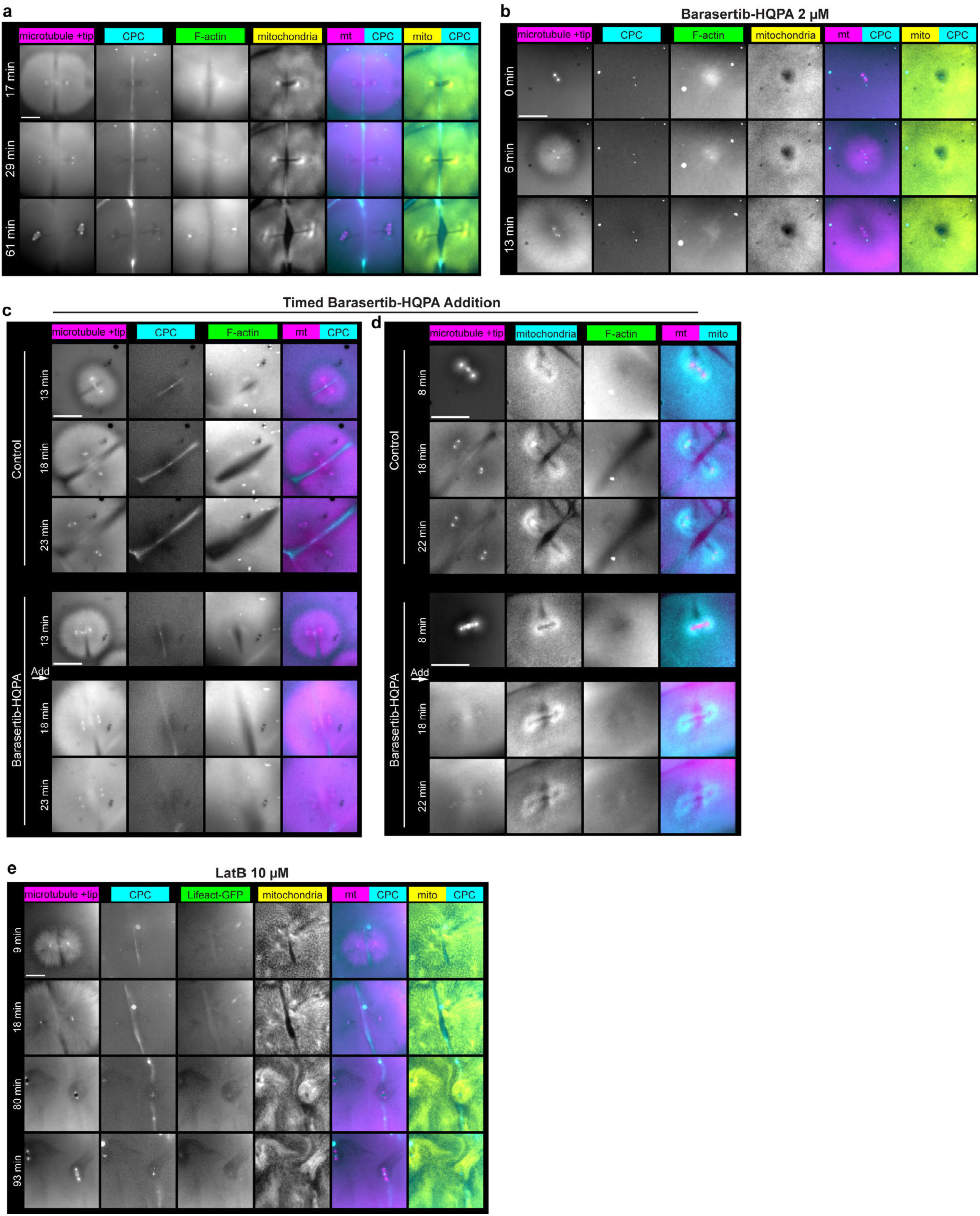
Dynamics and function of CPC in egg extract at the midplane, related to Fig. 3a**-c**. **a**, CPC localization co-imaged with microtubules, F-actin, and mitochondria, from late anaphase to the onset of the following mitosis. **b**, Mitotic dynamics of actin-intact cycling egg extract treated with 2 μM barasertib-HQPA. Note the lack of spindle assembly or centrosome separation. **c**, Bottom: timed local addition of barasertib-HQPA into cycling extract reaction with visualization of CPC, microtubules, and F-actin. As the extract reached anaphase, 0.2-0.3 μl of 0.5 mM barasertib-HQPA dissolved in egg extract was pipetted into the reaction adjacent to this pair of asters. Pipetting occurred at approximately 16 min. Top: control reaction without perturbation. **d**, Similar as **c**; bottom: local addition of barasertib-HQPA at approximately 11 min, visualizing mitochondria (MitoView Fix 640), microtubules, and F-actin. **e**, Dynamics of CPC and mitochondria (MitoView 405) in cycling egg extract where F-actin was depolymerized with 10 μM LatB. All scale bars, 200 μm. Time 0 = first frame with assembled spindles (and fully disassembled nuclei). Imaging probes include EB1-mApple (microtubule +tip), Lifeact-GFP (F-actin), anti-INCENP-AF647 (CPC) and MitoView 405 or MitoView Fix 640 (mitochondria).

**Extended Data Fig. 4:**
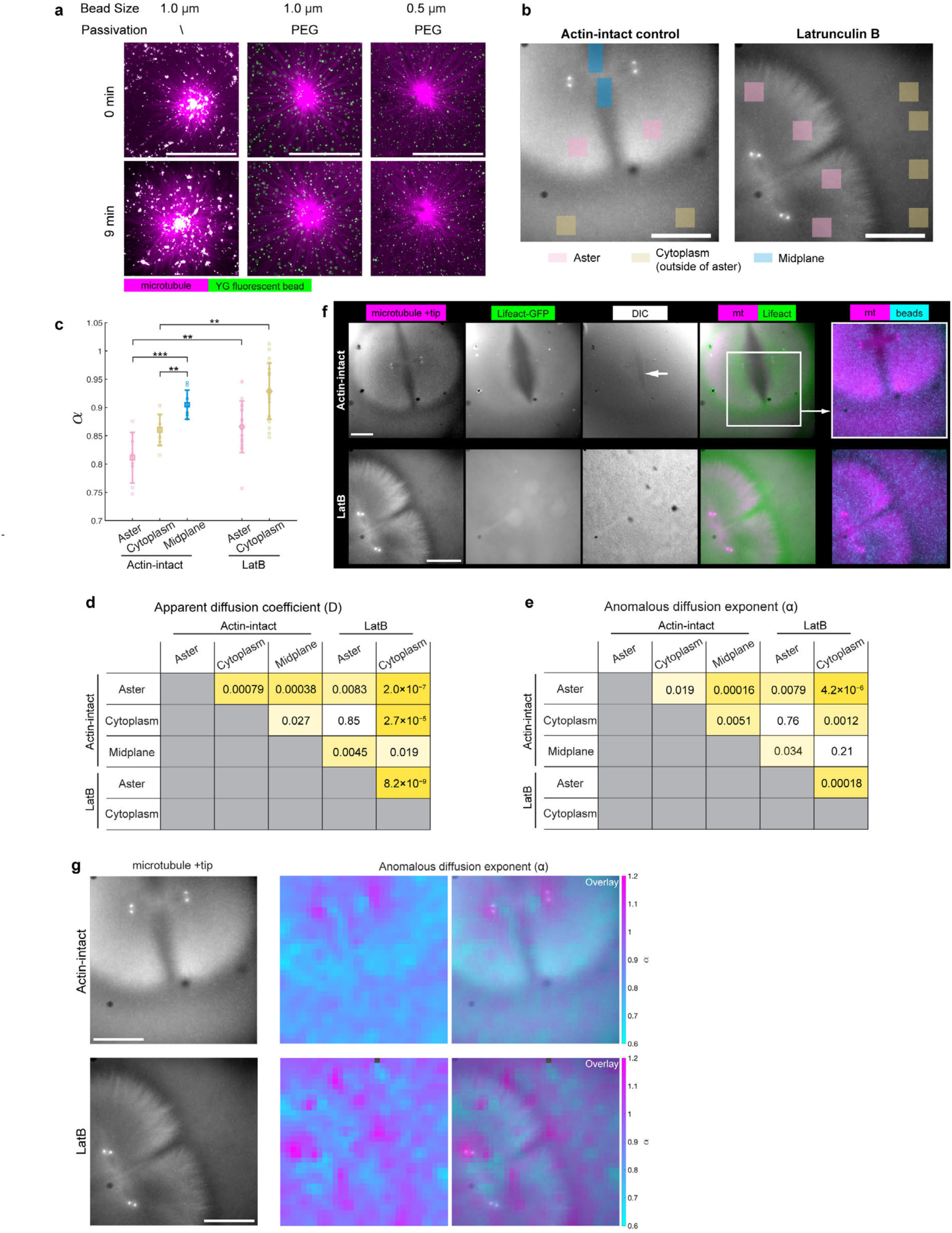
Characterization of cytoplasmic material properties, related to Fig. 3d**-h**. **a**, Dynamics of fluorescent beads, untreated and PEG-passivated, in interphase *Xenopus* egg extract prepared with CytoD. Note the strong aggregation and transport of untreated beads, versus the stable dispersion of PEG-passivated beads. **b**, Examples of sampled areas from different cytoplasmic regions in actin-intact and LatB-treated egg extracts (Fig. 3g and Extended Data Fig. 4c-e below). **c**, Anomalous diffusion coefficient (α) calculated from the same experiments described in Fig. 3g. *p < 0.05, **p<0.01, ***p<0.001, and ****p<0.0001 by Student’s t test. **d** and **e**, Tables of pairwise p values by Student’s t test, for D and α calculated from different cytoplasmic regions in actin-intact and LatB-treated egg extract (Fig. 3g and Extended Data Fig. 4c above). Shades of yellow correspond to significance levels, with darker yellow indicating more significant difference for the pair. No shading indicates no significant difference. **f**, Live images of microtubules (EB1-mApple), F-actin (Lifeact-GFP) and DIC images for actin-intact and LatB-treated egg extract, corresponding to Fig. 3h and Extended Data Fig. 4b. The high-diffusivity region at the midplane in Fig. 3h corresponds to the actin depletion zone with low contrast in DIC (white arrow). **g**, Local α calculated within overlapping 42x42 µm^2^ grids, locally averaged, and overlaid onto microtubule images, for actin-intact egg extract (top) and extract with 10 µM LatB (bottom). Related to Fig. 3h. All scale bar, 200 µm.

**Extended Data Fig. 5:**
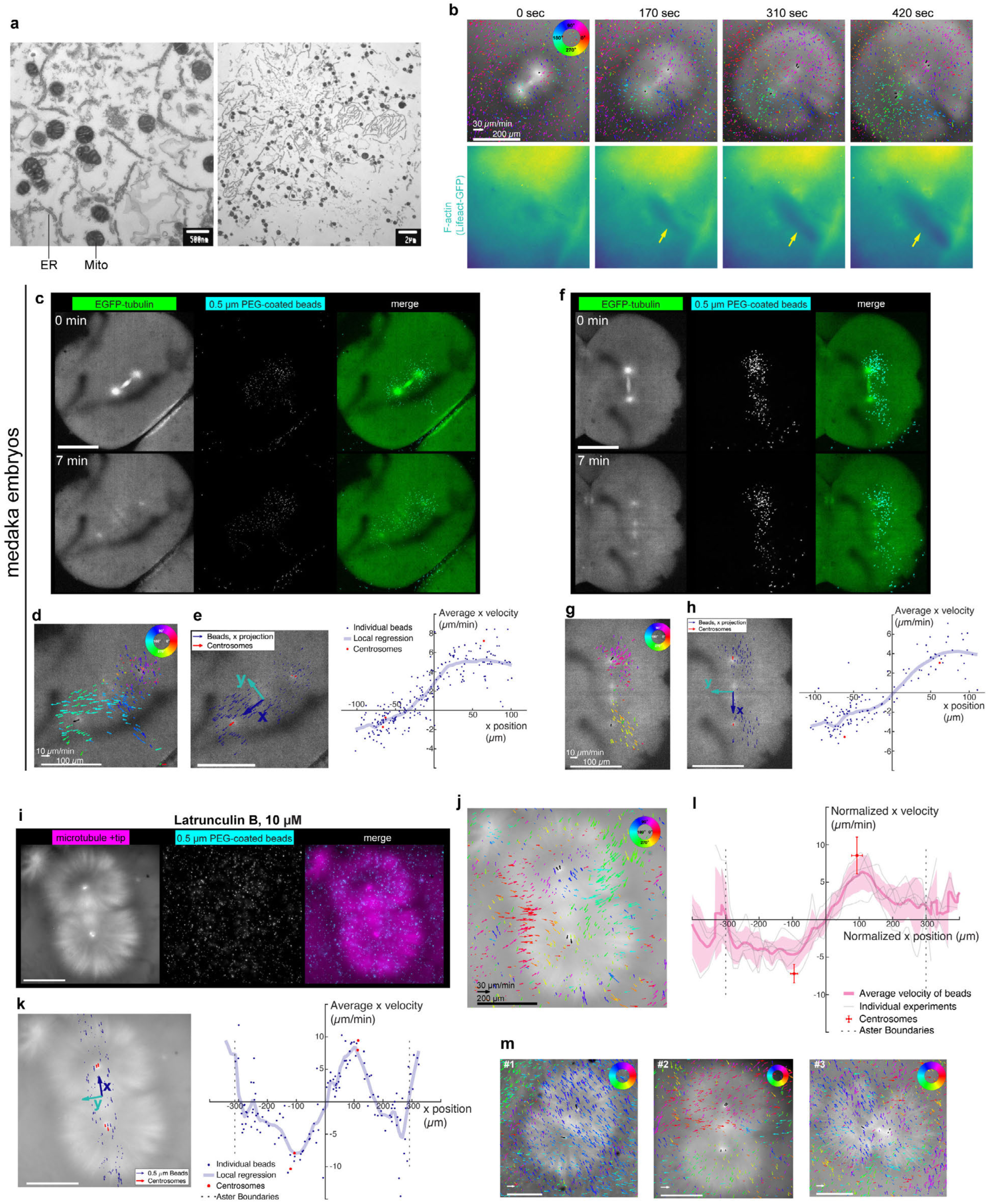
Tracking cytoplasmic flows with PEG-coated beads, related to Fig. 4. **a**, EM images of *Xenopus* egg extract, showing mitochondria, ER etc. Note the size of mitochondria is on the order of 0.5 μm. **b**, Cytoplasmic flows emerged as F-actin was progressively depleted along the midplane (highlighted by yellow arrows). **c** and **f**, Two medaka embryos with PEG-passivated beads injected, imaged at 2-cell to 4-cell stage. Live images of microtubules (EGFP-tubulin) and PEG-coated Fluoresbrite 641 beads were MaxIP from 11 Z-stacks with a step size of 2 µm. **d** and **g**, Velocity vectors from particle tracking of beads (colored arrows) and centrosomes (black arrows) overlaid on the microtubule image, averaged over 40 seconds. Velocity vectors of beads are color coded by orientation. **e** and **h**, Projection of velocity vectors in **d** and **g** onto the separation axis (designated as *x* axis) connecting daughter centrosomes, for centrosomes and beads within a 100-µm-wide band along the *x* axis. Projected *x* velocities were plotted against particle position along the x axis, and local regression with LOWESS was shown. **i**, Microtubule asters (EB1-mApple) and 0.5 µm PEG-passivated beads in 10 μM LatB-treated egg extract, imaged live during late anaphase. Images are MaxIP from 3 Z-stacks over 10 µm. **j**, Velocity vectors from particle tracking of beads (colored arrows) and centrosomes (black arrows) in **i** overlaid on the microtubule image, averaged over 40 seconds. Velocity vectors of beads are color coded by orientation. **k**, Velocity vectors in **j** within a 100-µm-wide band across centrosomes were projected onto the *x* axis. *x* components of particle velocity were plotted against particle position along the *x* axis, along with local regression with LOWESS. **l**, Ensemble profile of cytoplasmic velocity in LatB-treated extracts from 5 experiments prepared with 4 different egg extracts. Asters of similar radius (270 μm – 330 μm) were normalized to 300 μm for aggregated view; bead velocity was normalized so that bulk movement was set to 0. Gray lines show local regression of individual experiments; note the variation and fluctuation among experiments. The shaded curve represents mean ± SD of regression curves. **m**, Cytoplasmic velocity in LatB-treated cycling extract prior to normalization for 3 experiments in **l**, showing the variability among experiments. Scale bars in **c** - **h**, 100 µm. Linear scale bars in **i** - **m**, 200 μm; arrow scale bars in **m**, 30 μm/min.

**Extended Data Fig. 6:**
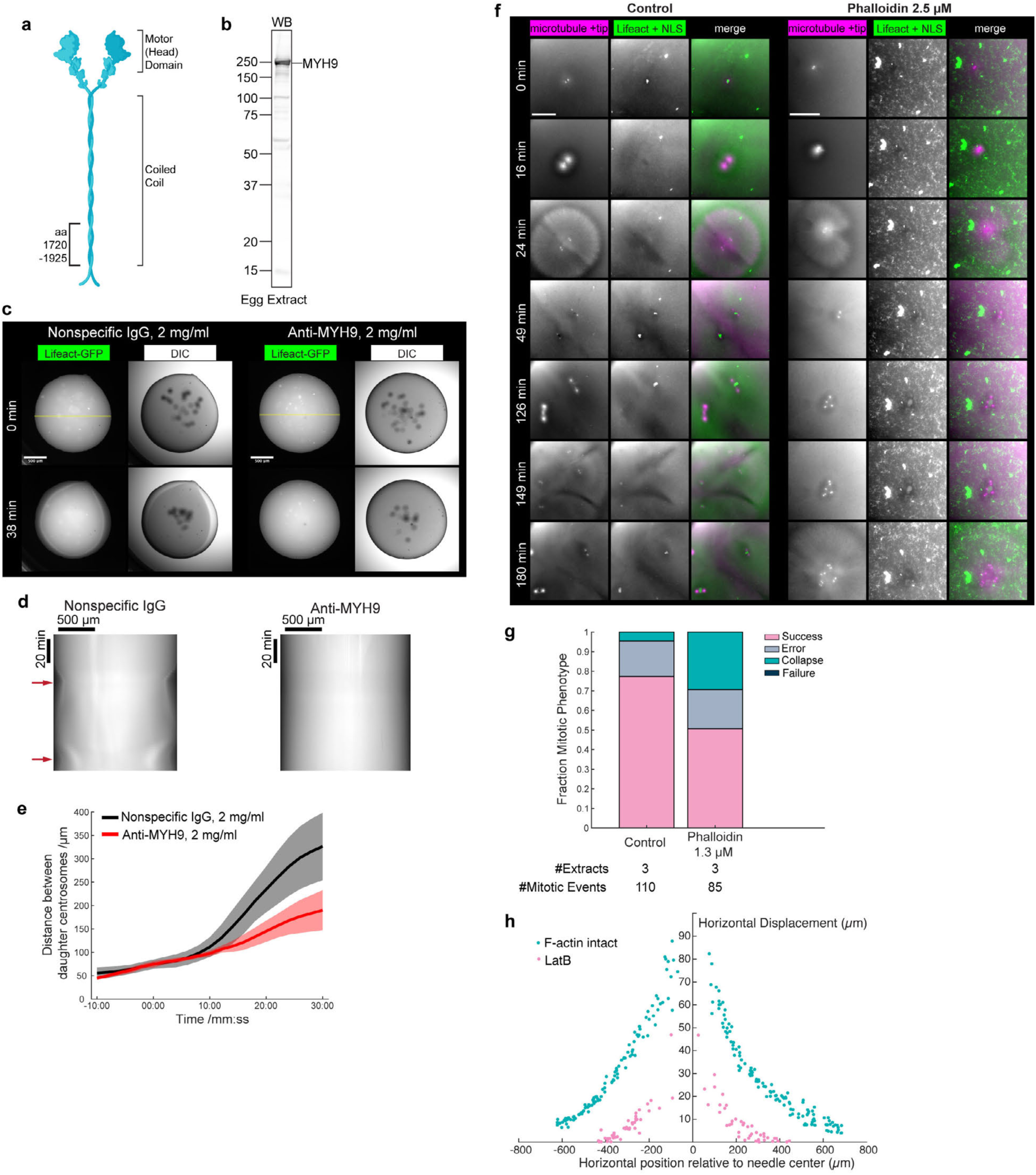
Myosin-II inhibition, F-actin stabilization, and bulk deformation. Related to Fig. 4 and 5. **a**, Diagram of non-muscle myosin II showing the coiled-coil domain in its heavy chain (MYH9) used as the antigen for raising inhibitory antibody. This C-terminal domain (amino acid 1720-1925) corresponds to the “assembly critical domain” (ACD) required for mini-filament assembly. **b**, The affinity-purified rabbit polyclonal antibody against MYH9 recognized a single band in *Xenopus* egg extract by Western blot (WB), consistent with the expected molecular weight of 226.8 kDa. **c**, Cycling egg extract was mixed with Lifeact-GFP and 2 mg/ml of either nonspecific rabbit IgG (as control) or MYH9 antibody. 1 μL droplets of each extract mixture were loaded under mineral oil and imaged live. Bulk contraction of actomyosin was seen with nonspecific rabbit IgG, but not with MYH9 antibody. **d**, Kymograph of bulk actomyosin dynamics in **c**, plotted along the diameter (20 pixel-wide lines indicated in yellow above) across the egg extract droplets over 112 min. Red arrows indicate bulk contraction during mitosis. **e**, Centrosome separation quantified for control and myosin-inhibited egg extract, as a repeat to Fig. 4g with a different egg extract from a different frog. n = 7 aster pairs in control and n = 10 in anti-MYH9-treated extract. Data are represented as mean ± SD. **f**, Cell cycle dynamics in control and 2.5 μM phalloidin-treated egg extract. Microtubules and centrosomes visualized with EB1-mApple; F-actin and nuclei visualized with Lifeact-GFP and GFP-NLS. Note the lack of bipolar spindles, sister asters, or nuclear/centrosomal separation with phalloidin treatment. Scale bars, 200 μm. **g**, Summary of mitotic outcomes among 3 pairs of control and 1.3 μM phalloidin-treated egg extract, corresponding to Fig. 4k and 4l. **h**, Horizontal displacement of beads along the path of needle movement (yellow shaded in Fig. 5b) plotted against horizontal position relative to needle center, for the experiments shown in Fig. 5b. Note the symmetrical displacement on both sides of the needle.

**Extended Data Fig. 7:**
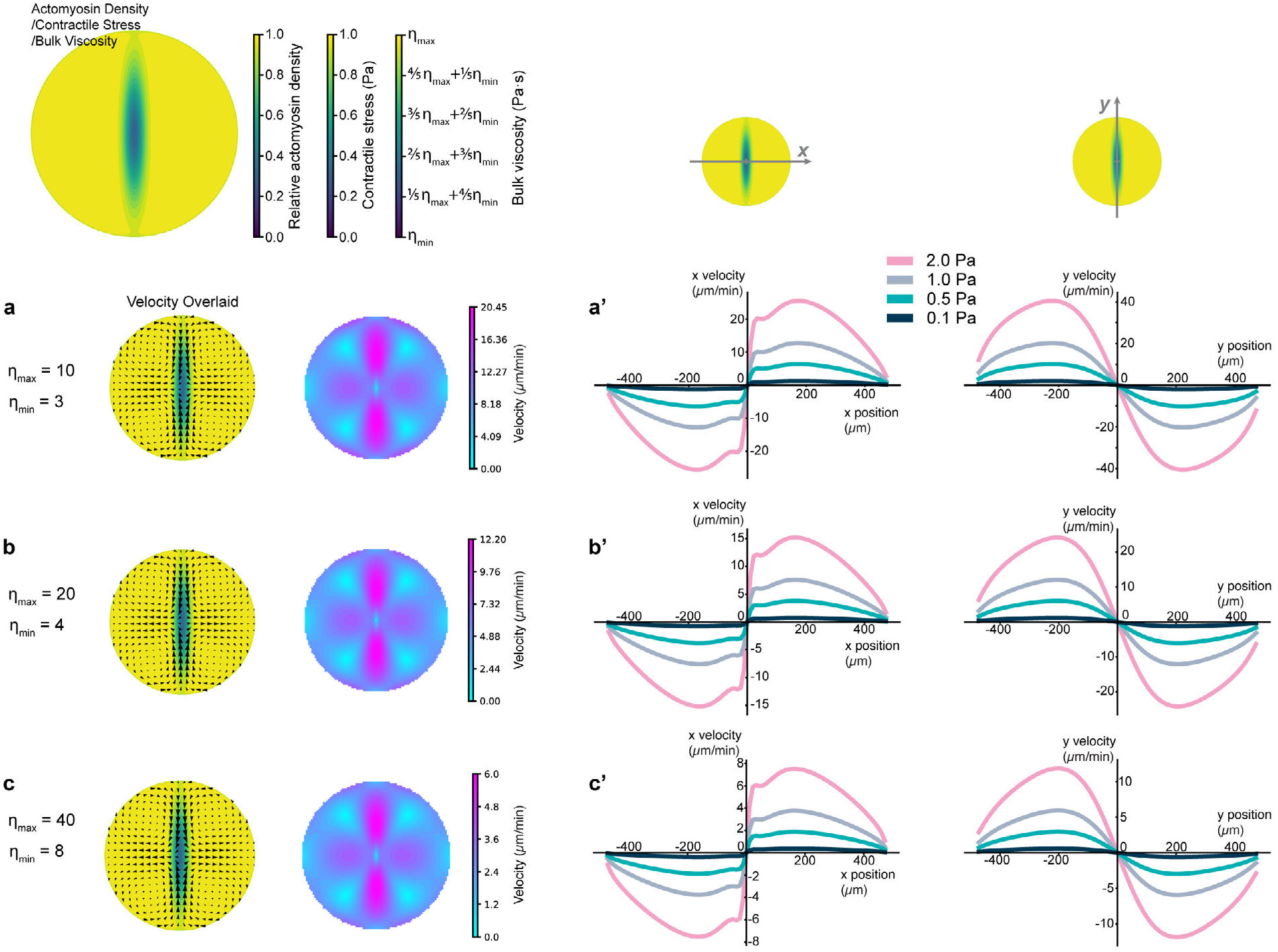
Simulation of cytoplasmic flows with varying parameters of bulk viscosity and contractile stress, related to Fig. 6. Consistent with Fig. 6b, a low-stress low-viscosity Gaussian ellipsoid simulated the actin depletion zone in a post-mitotic embryonic cell. Both contractile stress (0 – 1.0 Pa) and bulk viscosity (η*_min_*– η*_max_* Pa·s) were set to be linearly dependent on local actomyosin density. Diameter of the cell set as 1 mm. **a**, 2D simulation of cytoplasmic velocity from solving the Stokes equation with η*_max_* = 10 and η*_min_* = 3. **a’**, Cytoplasmic velocity along the *x* axis (y = 0) and along the *y* axis (x = 0), respectively, when varying the maximum contractile stress between 0.1 Pa and 2.0 Pa. **b** and **b’**, Same as **a** and **a’** except η*_max_* = 20 and η*_min_* = 4. Here same parameters as in Fig. 6 were re-used for side-by-side comparison. **c** and **c’**, Same as **a** and **a’** except η*_max_* = 40 and η*_min_* = 8.

**Supplementary Video 1**, related to Fig. 1. Dynamics of microtubules (Tau-mCherry) and nuclei (GFP-NLS) in actin-intact cycling egg extract, with DIC images showing cytoplasmic compartmentalization. MaxIP from 2 z-stacks.

**Supplementary Video 2**, related to Fig. 1. Microtubule (Tau-mCherry) dynamics during continuous cell cycle progression over 4 hours in actin-intact cycling egg extract.

**Supplementary Video 3**, related to Fig. 2. Partitioning of F-actin (Lifeact-GFP) and acidic vesicles (LysoTracker Deep Red) along the midplane, co-imaged with microtubule plus ends (EB1-mApple) in actin-intact cycling egg extract. MaxIP from 2 z-stacks.

**Supplementary Video 4**, related to Fig. 2 and Extended Data Fig. 2. Dynamics of acidic vesicles (LysoTracker Deep Red) in cycling egg extract treated with 10 μM LatB to depolymerize F-actin, co-imaged with microtubule plus ends (EB1-mApple) and Lifeact-GFP.

**Supplementary Video 5**, related to Fig. 3. Dynamics of CPC (anti-INCENP) from metaphase to interphase in actin-intact cycling egg extract, co-imaged with microtubule plus ends (EB1-mApple), F-actin (Lifeact-GFP), and mitochondria (MitoView 405).

**Supplementary Video 6**, related to Fig. 4. Cytoplasmic flows during late anaphase in actin-intact cycling egg extract. Left: timelapse images of PEG-passivated Fluoresbrite 641 beads 0.5 μm and microtubule plus ends (EB1-mApple). Right: bead velocity averaged over 40 sec, color-coded by orientation, and overlaid on EB1-mApple images. MaxIP from 3 z-stacks.

**Supplementary Video 7**, related to Fig. 4 and Extended Data Fig. 4. Cytoplasmic flows in a medaka embryo at 2-cell to 4-cell stage. Left: timelapse images of PEG-passivated Fluoresbrite 641 beads 0.5 μm and microtubules (EGFP-tubulin). Right: bead velocity averaged over 40 sec, color-coded by orientation, and overlaid on EGFP-tubulin images. MaxIP from 11 z-stacks.

**Supplementary Video 8**, related to Fig. 5. Stretching microtubule asters (Tau-mCherry) with embedded 1 μm PEG-passivated Fluoresbrite YG beads in actin-intact cycling egg extract with a needle.

**Supplementary Video 9**, related to Fig. 5. Stretching microtubule asters (Tau-mCherry) with embedded 1 μm PEG-passivated Fluoresbrite YG beads in cycling egg extract treated with 10 μM LatB.

**Supplementary Video 10**, related to Fig. 6. Simulation of cytoplasmic flows with the emerging actin depletion zone.

## Notes

### Competing Interest Statement

The authors have declared no competing interest.

